# Non-canonical H3K79me2-dependent pathways promote the survival of MLL-rearranged leukemia

**DOI:** 10.1101/2020.12.04.411215

**Authors:** William F. Richter, Rohan N. Shah, Alexander J. Ruthenburg

## Abstract

MLL-rearranged leukemia depends on H3K79 methylation. Depletion of this transcriptionally-activating mark by DOT1L deletion or high concentrations of the inhibitor pinometostat downregulates *HOXA9* and *MEIS1*, and consequently reduces leukemia survival. Yet some MLL-rearranged leukemias are inexplicably susceptible to low-dose pinometostat, far below concentrations that downregulate this canonical proliferation pathway. In this context, we define alternative proliferation pathways that more directly derive from H3K79me2 loss. By ICeChIP-seq, H3K79me2 is markedly depleted at pinometostat-downregulated and MLL-fusion targets, with paradoxical increases of H3K4me3 and loss of H3K27me3. Although downregulation of polycomb components accounts for some of the proliferation defect, transcriptional downregulation of FLT3 is the major pathway. Loss-of-FLT3-function recapitulates the cytotoxicity and gene expression consequences of low-dose pinometostat, whereas overexpression of constitutively active *STAT5A*, a target of FLT3-ITD-signalling, largely rescues these defects. This pathway also depends on MLL1, indicating combinations of DOT1L, MLL1 and FLT3 inhibitors should be explored for treating *FLT3-*mutant leukemia.

## Introduction

MLL1-rearrangements (MLL-r) account for ~10% of all leukemia cases and are especially prominent in infants (70-80%) and, lacking an effective standard of care, bear a very poor prognosis^1–5^. A growing body of evidence suggests that MLL-rearrangements rely on additional mutations to cause leukemia. Leukemia patients with MLL-fusions often have additional mutations that affect growth signaling pathways^6–8^ and MLL-fusions in mouse models cause leukemias with longer-than-expected latencies, suggesting that additional mutations are required for full progression^9–11^. Yet few studies have examined the genetic context of MLL-fusion proteins and how additional lesions may cooperate to promote disease at the molecular level.

*MLL1* (Mixed Lineage Leukemia protein, also known as *KMT2A*) is a histone H3 lysine methyltransferase involved in regulating *HOX* gene expression during development and normal hematopoiesis^12^. Translocations of MLL1 fuse its amino terminus to the carboxy-terminus of a growing list of over 130 different fusion partners^13^. Although these MLL-fusions lack methyltransferase activity, a functional copy of the MLL1 gene is necessary to target and hypermethylate H3K4 at MLL-fusion target genes to induce leukemogenesis^14–16^. In more than 75% of acute myeloid leukemia (AML) cases and > 90% of acute lymphoblastic leukemia (ALL) cases involving MLL translocations, the MLL-fusion partner is one of 7 members of the transcriptional elongation complex, most commonly, AF9 and AF4 respectively^17^. These fusion partners aberrantly recruit DOT1L, the sole histone H3 lysine 79 methyltransferase to MLL1 target genes including the HOXA gene cluster ^18–20^. By mechanisms that remain unclear, DOT1L-mediated hypermethylation of H3K79 promotes expression of MLL-fusion targets^14,21–24^, establishing an expression profile with a surprising degree of target gene overlap across different MLL-fusions^25^. Ablation of H3K79 methylation through knockout or pharmacological targeting of *DOT1L* abrogates the MLL-fusion target gene expression profile, selectively induces apoptosis and differentiation of leukemia cells in culture and dramatically extends the survival of mice in xenograft experiments^21,26^.

Viral co-transduction of the MLL-AF4 targets^27^ *HOXA9* and *MEIS1* is sufficient to cause acute leukemia in mouse bone marrow progenitors, arguing that these transcription factors represent a major etiologic pathway in MLL-r leukemia^10,28–31^. However, exogenous expression of *MLL-AF9* in mice requires a long latency period (4-9 months) and chemotherapy induced MLL-translocations cause disease 3-5 years after treatment, suggesting that additional mutations are required for leukemagenesis^10,32^. In the prevailing model, MLL-fusions recruit DOT1L to hypermethylate and activate expression of *MEIS1* and *HOXA9* (Figure 1A)^19,21,22,33,34^. However, the genetic manipulations used to define this paradigm may have missed more subtle and graded effects afforded by kinetically-staged antagonism with highly-specific small-molecule inhibitors. Therefore, to better understand the direct effects of H3K79me2 in several MLL-r cell lines we employed pharmacologic inhibition of DOT1L methyltransferase activity.

**Figure 1.**
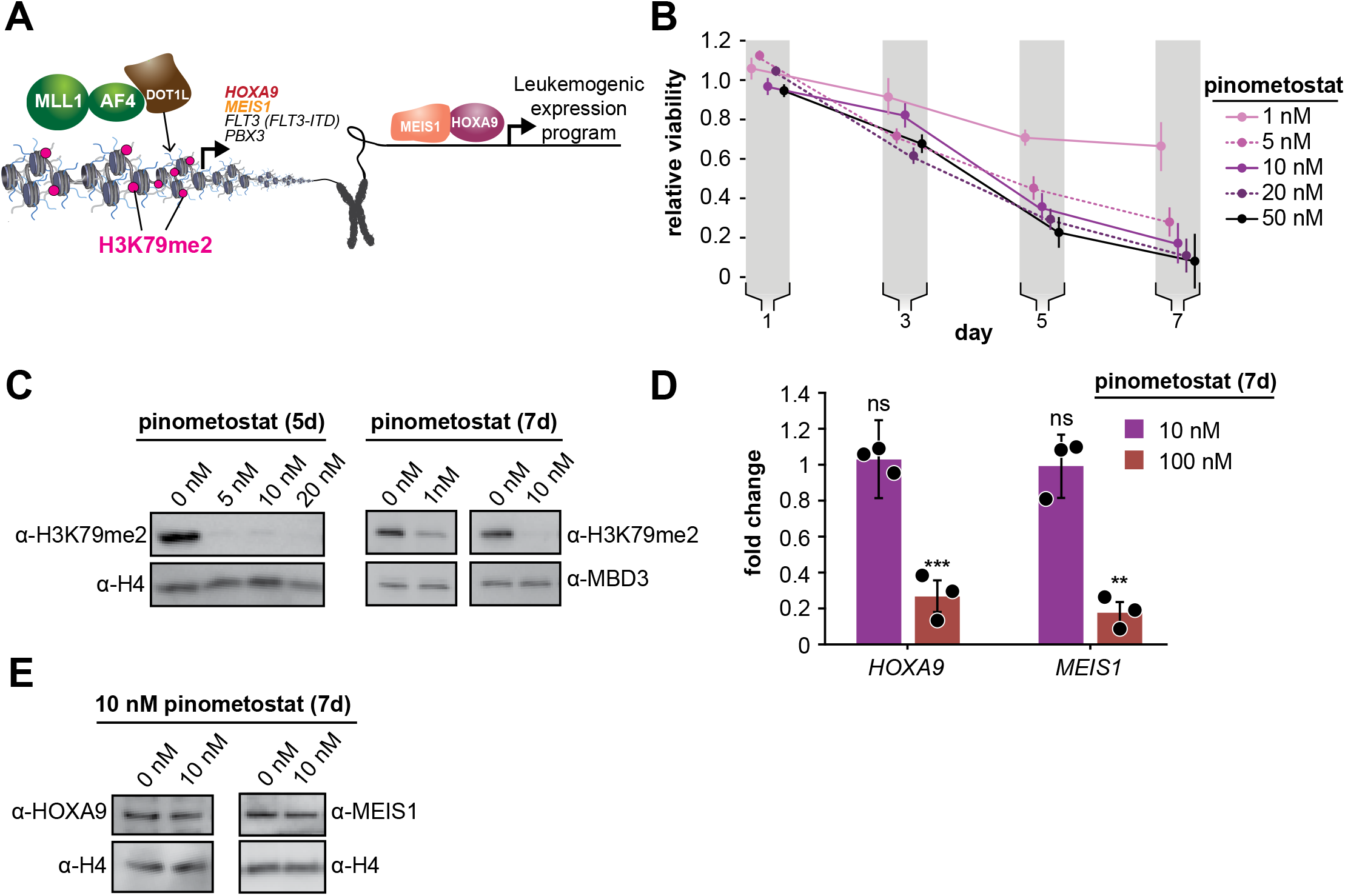
Low doses of DOT1L inhibitor ablate bulk H3K79me2 and curtail MV4;11 proliferation without impacting expression of canonical target genes. **A.** Conventional model depicting how DOT1L methyltransferase activity activates transcription of key proliferative oncogenic transcription factors^19,21,22,25,27,30^. **B.** Proliferation assay of MV4;11 cells treated with the indicated concentrations of the DOT1L inhibitor pinometostat (EPZ5676). Cell viability was assayed every two days, starting one day after treatment commenced using the CellTiter Glo 2.0 reagent. Relative cell viability is presented as the mean fraction of pinometostat versus cells treated with the equivalent volume of DMSO from three independent experiments ± S.E.M. **C.** Western blots for H3K79me2 with H4 or MBD3 loading controls in MV4;11 cells treated with 1 to 20 nM pinometostat for 5 or 7 days. **D.** RT-qPCR analysis of *HOXA9* and *MEIS1* expression fold-change in MV4;11 cells treated with 10 or 100 nM pinometostat for 7 days. Results are shown as mean ± S.E.M. of three independent experiments. Student’s t-test (ns *p* > 0.05, ** *p* ≤ 0.01, *** *p* ≤ 0.001). **E.** Western blot of HOXA9 and MEIS1 with H4 as a loading control from MV4;11 cells treated with 10 nM pinometostat for 7 days.

Pinometostat (EPZ5676), a highly specific DOT1L inhibitor^33,35,36^ displays 37,000-fold selectivity over its closest related paralogs and a host of other lysine and arginine methyltransferases^6^. Interestingly, several cell lines that all have the *MLL-AF4* translocation display pinometostat sensitivities that differ by nearly 3 orders of magnitude^26^. One of these lines (MV4;11) displays a pinometostat IC50 for proliferation that is 20 times lower than the IC50 for *HOXA9* and *MEIS1* expression^26^, suggesting that these drivers of leukemogenesis, though downregulated at higher concentrations (1 μM)^26^, may not contribute to cell-type specific effects at lower concentrations.

We sought to understand low-dose pinometostat effects by treating a variety of MLL-r cell lines with a concentration that reduces proliferation in only a subset, with MLL-r cell lines harboring *FLT3-ITD* mutations being the most susceptible. Under these conditions, *HOXA9* and *MEIS1* expression remain unaffected, presenting a clear exception to the existing paradigm, but found thousands of other differentially expressed genes, including the *PBX3* and *FLT3* oncogenes. Capitalizing on the sensitivity of internally calibrated ChIP-seq (ICeChIP-seq)^37,38^, we observed larger reductions in H3K79me2 density at a subset of MLL-AF4 targets, a genome-wide reduction in H3K27me3 and stark H3K4me3 increases at transcription start sites.

Remarkably, we could nearly completely rescue not only pinometostat-but also MLL1 inhibitor-induced effects on proliferation and apoptosis through expression of a constitutively active form of the downstream FLT3-ITD target *STAT5A* (*STAT5A-CA*), arguing that disruptions to this pathway represent the main source of toxicity from low-dose DOT1L inhibition. In addition, DOT1L inhibition also downregulated the *EZH2* and *EED* components of the PRC2 complex, likely accounting for global reductions in H3K27me3 and imparting modest, but distinct effects on proliferation and a correspondingly moderate proliferation rescue from EZH2 overexpression. Collectively, our data argue that the FLT3-ITD signaling and PRC2 pathways, are more sensitive to disruptions of MLL-fusion-mediated gene activation than the canonical oncogenic drivers in MLL-r, *FLT3^ITD^* leukemias, defining a new molecular understanding of how MLL-fusions cooperate with other oncogenic factors to induce leukemia.

## Results

### MLL-r leukemia is sensitive to DOT1L inhibitor via a non-canonical pathway

Leukemias harboring MLL-rearrangements are uniquely susceptible to DOT1L inhibition and MV4;11, a biphenotypic leukemia cell line harboring an *MLL-AF4* translocation, is one of the most sensitive^26^. To determine the basis of this susceptibility we systematically examined how low-dose regimes of pinometostat affect proliferation and global H3K79me2 levels in cells treated for 7 days with 1-50 nM pinometostat. This range of concentrations encompasses the previously determined MV4;11 proliferation IC50 (3.5 nM) but is well below the 1 μM or higher typically used in published investigations of the effects of H3K79me ablation^26,39,40^. Consonant with previous findings^26^, pinometostat concentrations as low as 1 nM significantly reduce global levels of H3K79me2 and cause a 30% ± 10% reduction in MV4;11 proliferation, while 10 nM inhibitor reduced cell proliferation by 80% ± 10% (Figure 1B and C). Notably, after treating MV4;11 cells with 10 nM inhibitor for 7 days we observed no discernable effect on the expression of *HOXA9* and *MEIS1* (Figure 1D and E), despite the emphasis on these genes as the critical mediators of DOT1L’s effects in MLL-r leukemia^19,21,22,33,34^. Consistent with prior observations^26^, a much higher dose of 100 nM pinometostat significantly downregulates both *HOXA9* and *MEIS1* expression (Figure 1D).

### DOT1L inhibition at low concentrations downregulates leukemic oncogenes

With the extant model^19,21,22, 33,34^ unable to explain reductions in proliferation caused by the DOT1L inhibitor in this concentration regime, we reasoned that the expression of other genes crucial to the survival of these cells are likely affected. To define these genes, we performed RNA-seq in MV4;11 cells that had been treated with 10 nM pinometostat for 7 days and observed that 1916 genes were downregulated and 2007 genes were upregulated (Figure 2A) relative to a DMSO treated control. To account for any handling biases, we included 4 RNA “spike-in” controls and found no significant differences in read counts between treatment groups (Figure S1A). The downregulated genes significantly overlap with MLL-AF4 targets identified by Kerry et al. by ChIP-seq in MV4;11 cells^20^ (Figure 2B). Relative to prior high-dose (3 μM) treatment with a compound structurally related to pinometostat in MV4;11 cells, the numbers of differentially expressed genes are similar, and there is marked overlap between the sets, particularly the downregulated cohort^33^ (Figure 2C and S1B). Consistent with our RT-qPCR measurements, *HOXA9* was unaltered in its expression (Figure S1C) and *MEIS1* displayed extremely modest mRNA reduction (20%) not observed by RT-qPCR and not reflected in apparent protein levels (Figure 1D-E). Of the other *HOXA* cluster genes only *HOXA11* and *HOXA13* exhibited expression changes with a 1.7-fold decrease and 2.5-fold increase, respectively (Figure S1C).

**Figure 2.**
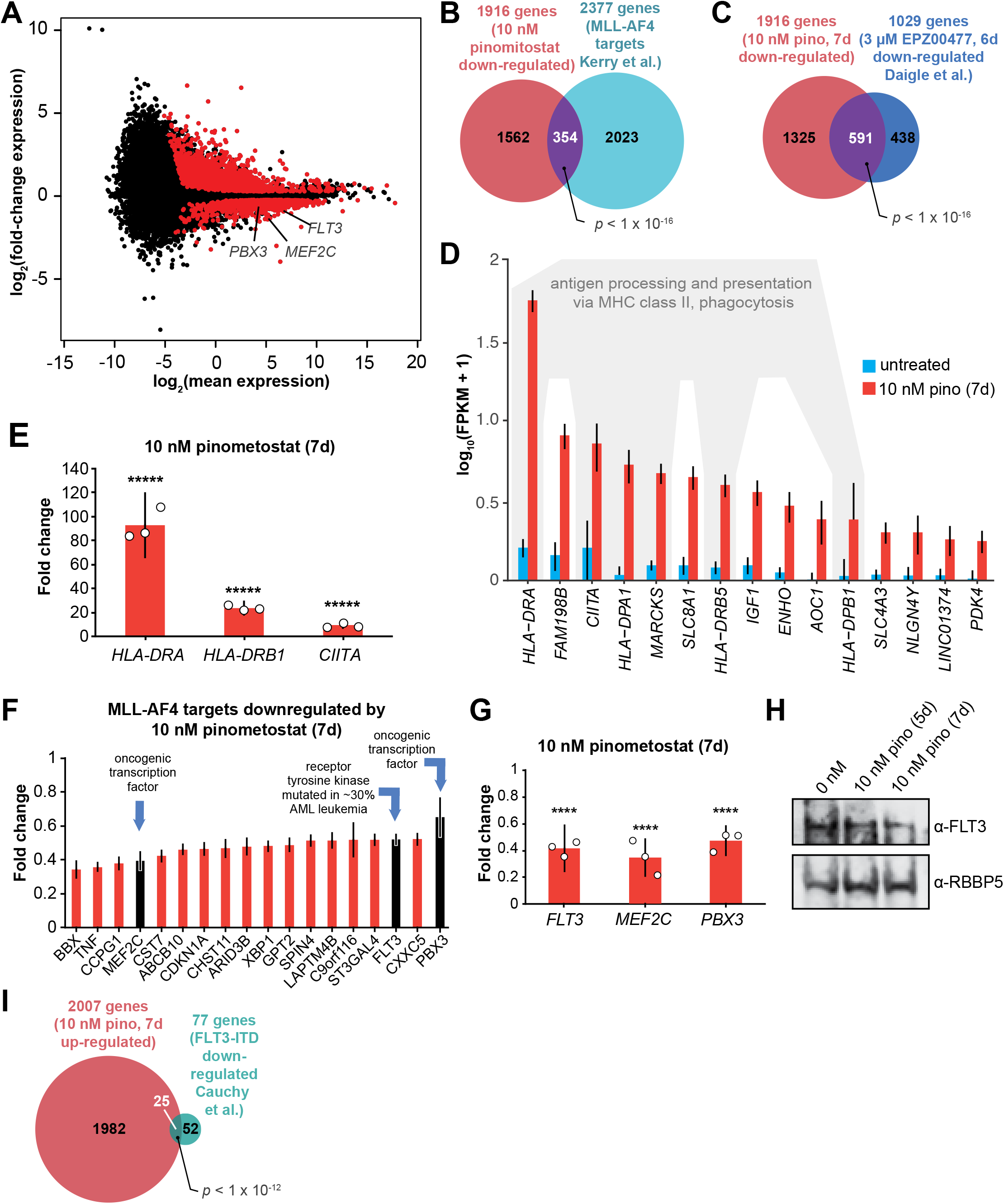
DOT1L inhibition downregulates a subset of MLL-AF4 targets including the leukemic oncogene FLT3. **A.** MA plot showing genes differentially expressed in MV4;11 cells treated with 10 nM pinometostat or DMSO 7 days as log_2_-mean of expression (FPKM) of the DMSO and pinometostat treated samples versus the log_2_-fold change of the mean normalized pinometostat versus DMSO treated FPKM for three independent replicates. Red represents genes that meet the significance threshold, with an FDR-adjusted *p* ≤ 0.5. **B.** Venn diagram depicting overlapping genes between those downregulated by 10 nM pinometostat and MV4;11 MLL-AF4 targets identified by Kerry et al.^20^, *p*-value computed by two-tailed Fisher Exact test. **C.** Venn diagram displaying the overlap between genes downregulated in MV4;11 cells by 10 nM pinometostat treatment (7 days) and treatment with 3 μM of the pinometostat-related compound EPZ004777 for 6 days^33^. *p*-value computed by two-tailed Fisher Exact test. **D.** Bar plot depicting upregulated genes with the highest fold changes from RNA-seq analysis of 3 independent experiments of DMSO-(blue) or pinometostat-treated (red) MV4;11 cells with uncertainty presented as the standard deviation computed by CuffDiff^116^ with immune response genes outlined in gray. **E.** RT-qPCR analysis showing the fold-change for *HLA-DRA, HLA-DRB1* and *CIITA* gene expression in MV4;11 cells ± 10 nM pinometostat treatment for 7 days. Results are shown as mean ± S.E.M. of three independent experiments. Student’s t-test (\*\*\*\*\**p* ≤ 0.00001) **F.** Bar plot depicting the top pinometostat-downregulated genes from the RNA-seq analysis that are previously described MLL-AF4 targets^22^ including the oncogenes *MEF2C, FLT3* and *PBX3*. **G.** RT-qPCR analysis of *MEF2C, FLT3* and *PBX3* expression in MV4;11 cells ± with 10 nM pinometostat for 7 days. Results are displayed as mean fold-change ± S.E.M. of three independent experiments; Student’s t-test (**** *p* ≤ 0.0001). **H.** Western blot for FLT3 with RBBP5 as loading control in MV4;11 cells treated with 10 nM pinometostat for 5 or 7 days. **I.** Venn diagram displaying the overlap between genes upregulated in MV4;11 cells by 10 nM pinometostat treatment (7 days) and genes downregulated in leukemic cells from patients with *FLT3-ITD* vs normal *FLT3* karyotypically normal AML^61^. *p*-value computed by two-tailed Fisher Exact test.

Although H3K79me2 is considered transcriptionally activating, the upregulated genes had much larger expression fold-changes. 906 genes were upregulated at least 2-fold (and some > 80-fold), while only 86 genes were downregulated ≥ 2-fold (Figure 2A). The list of upregulated transcripts include MHC class II and innate immune response genes (Figure 2D). We confirmed the expression increases of *CIITA* (the master regulator of interferon-inducible MHC class II genes), and the MHC class II genes *HLA-DRA* and *HLA-DRB1* by RT-qPCR (Figure 2E). Gene ontology analysis of the upregulated genes indicated enrichment for “immune response” and “interferon-gamma signaling pathway” (Figure S2A)^41,42^. Despite there being no discernable effect on interferon-gamma (*IFNG*) expression in the RNA-seq analysis (Figure S2B), marked activation of IFN-γ-inducible genes is apparent. We hypothesize that this may be due to perturbations to signaling effectors of the IFN-γ pathway which includes the STAT family of transcription factors that are often aberrantly expressed in leukemia and other cancers^43–45^. The activation of so many genes involved in antigen processing and presentation as well as macrophage cell surface markers (Figure S2C) may indicate that these cells are undergoing differentiation towards a more macrophage-like state, consistent with apparent differentiation observed in other DOT1L loss-of-function paradigms^21,33^. By Gene Set Enrichment Analysis (GSEA)^46,47^ the set of differentially expressed genes were enriched for hematopoietic differentiation factors and anticorrelated with hematopoietic progenitor expression signatures (Figure S2D). Notably, the cytokine receptors CSF1R and CSF3R, critical signaling inducers of hematopoietic differentiation, were upregulated (Figure S2E)^48,49^.

Among the most downregulated genes were many MLL-AF4 target genes^20,22,50^ including the oncogene FMS-Like Tyrosine Kinase 3 (*FLT3*), the protooncogene Myocyte Enhancer Factor 2C (*MEF2C*) and Pre-B-cell leukemia homeobox 3 (*PBX3)* (Figure 2F). These genes all have previously described roles in the development of MLL-rearranged leukemias^23,51–53^. FLT3 is a receptor tyrosine kinase that regulates proliferation and cell survival via STAT and other signaling pathways. Mutations that constitutively activate FLT3 by internal tandem duplication of its juxtamembrane domain (FLT3-ITD) or point mutations within its kinase domain collectively represent the most frequently occuring genetic lesions in acute myeloid leukemia^51,54,55^. MV4;11 cells are homozygous for the *FLT3-ITD* mutation and highly sensitive to FLT3 inhibition^8,56^. The transcription factor *MEF2C* cooperates with *SOX4* to induce leukemogenesis in mouse models and *MLL-AF9*-expressing hematopoietic progenitors to promote colony formation^52,57^. PBX3 is a transcription factor that acts to stabilize both HOXA9 and MEIS1 localization at a subset of target genes and coexpression of either oncogene with *PBX3* can cause leukemogenesis^53,58,59^. We verified the reductions in *FLT3, MEF2C* and *PBX3* expression with pinometostat by RT-qPCR and examined FLT3 protein levels by Western blot (Figure 2G-H).

We wondered if downregulation of one or more of these genes could be responsible for the reductions in cell proliferation from low-dose pinometostat treatment. Using previously published datasets of MEF2C and FLT3-regulated genes, we first looked at the expression of 15 genes that were downregulated by MEF2C knockout in mouse hematopoietic progenitors^60^. Of these genes, only FLT3 was downregulated in our pinometostat-treated cells. Because the expression of nearly all of the set of MEF2C-regulated genes was unaffected in our analysis we moved our focus to FLT3. Previous work by Cauchy et al. identified 138 genes significantly upregulated in karyotypically normal *FLT3-ITD+* AML compared to WT *FLT3* AML patient samples^61^. A comparison of those FLT3-ITD-upregulated genes to our pinomeostat downregulated genes yielded a small but significant overlap (Figure S2F). We saw a more pronounced overlap between genes downregulated in FLT3-ITD+ patient samples and those upregulated by pinometostat, including 10 MHC class II receptors (Figure 2I). *PBX3* is the only MLL-AF4 target upregulated in the *FLT3-ITD* samples, suggesting it could be a crucial convergence point of the MLL-AF4 and FLT3-ITD pathways. Collectively, these data suggest that FLT3-ITD may represent an important pathway through which DOT1L inhibition reduces leukemia cell survival. Before delving further into the delineation of the responsible molecular pathways, we first sought to quantitatively define the consequences of low dose DOT1L inhibition on the distribution of the H3K79me2 mark and its causal connection to these gene expression-level changes.

### MLL-AF4 targets downregulated by low dose DOT1L inhibition are highly enriched for H3K79me2

Despite extensive global reductions in H3K79me2 levels, only a subset of MLL-AF4 targets were downregulated by 10 nM pinometostat, necessitating more nuanced measurement of the mark, particularly MLL-AF4 target genes. The current model, that MLL-AF4 recruits DOT1L to target genes resulting in aberrantly high levels of H3K79me2 and transcriptional activation^21,22,26^, has not been rigorously examined by quantitative methods that would be sensitive to small changes. Indeed, the limitations of conventional ChIP-seq preclude unambiguous quantitative analyses for direct comparisons of histone modifications upon global depletion^37,62^. To circumvent these problems, we used ICeChIP-seq, a form of native ChIP that uses barcoded internal-standard modified nucleosomes to permit direct quantitative comparison of histone modification density (HMD) at high resolution across samples^37,38,63^.

With ICeChIP we were able to measure a positive correlation (R^2^ = 0.53) between transcript abundance and H3K79me2 levels in MV4;11 cells (Figure 3A), consistent with the speculated role for H3K79me2 in transcriptional activation^19,21,24,33^. However, only 30 of the 250 most highly-expressed genes, including only 3 MLL-AF4 targets, were downregulated by 10 nM pinometostat treatment, suggesting that H3K79me2 is not necessary to maintain high levels of gene expression at all sites where it is enriched. The genes that were downregulated by 10 nM pinomeostat had higher H3K79me2 levels compared to upregulated genes or all expressed genes, rivalling the most highly expressed genes (Figure 3B). Although previous conventional ChIP-seq measurements observed enrichment of H3K79me2 at MLL-fusion target genes^21,22^, our ICe-ChIP-seq analysis revealed equivalent average density at MLL-AF4 targets and 250 most highly expressed genes (Figure 3B). Given that only 12 MLL-AF4 targets are included in that highly expressed gene list, this higher H3K79me2 density is likely due to very efficient recruitment of DOT1L by MLL-AF4 rather than deposition via the transcriptional apparatus^64,65^. Interestingly, the subset of MLL-AF4 targets that are downregulated by 10 nM pinometostat exhibit still higher levels of H3K79me2 than even MLL-AF4 targets as a whole and appear to be more dependent on H3K79me2 for their expression (Figure 3A and B). The only other group of genes analyzed with comparable peak H3K79me2 levels were “MLL-spreading” genes which display a binding profile that stretches further downstream into the gene body^20^.

**Figure 3.**
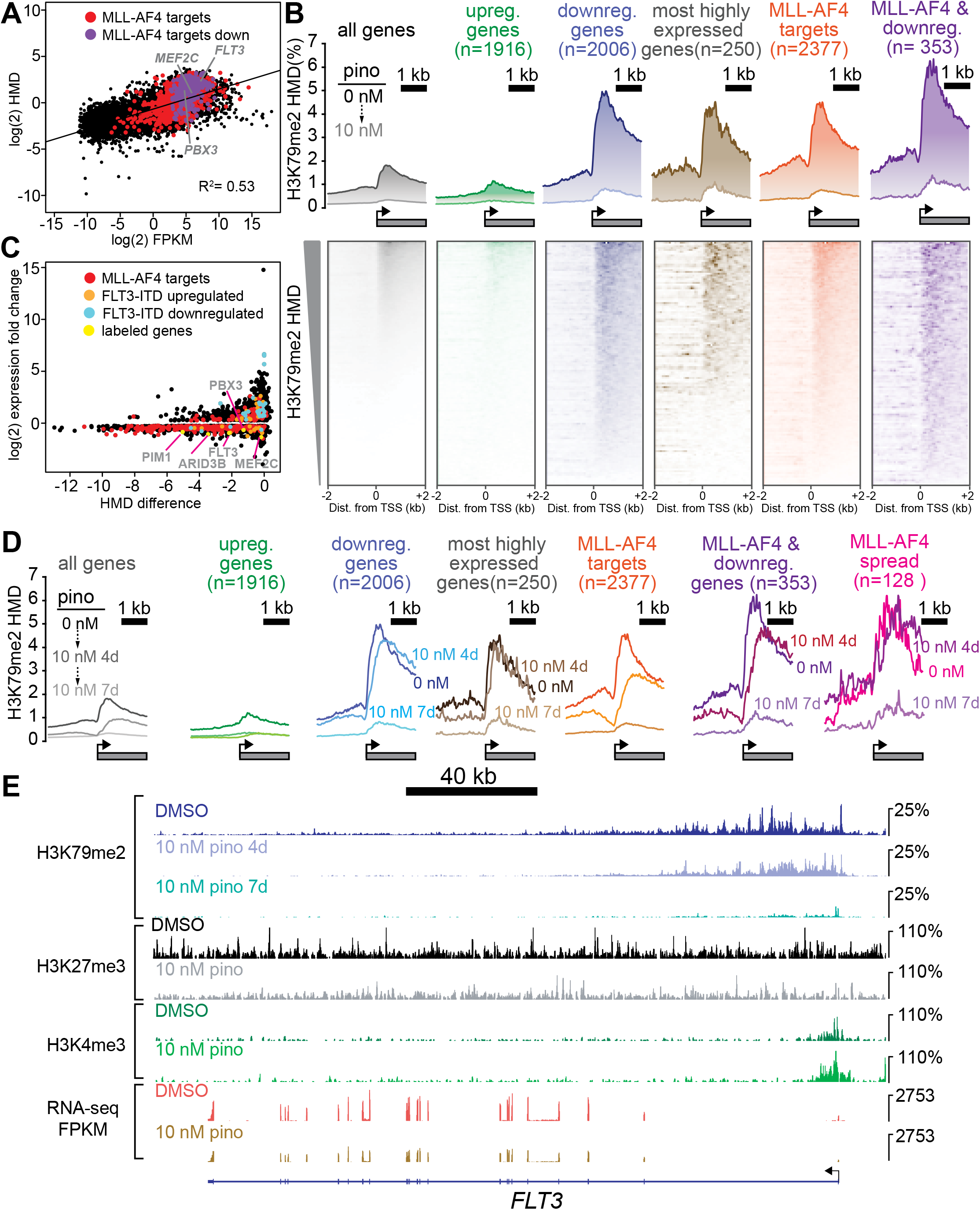
Low dose DOT1L inhibition disrupts H3K79me2 with more pronounced effects on downregulated MLL-AF4 targets. **A.** Scatterplot of the mean normalized log_2-_FPKM (three independent replicates) of genes expressed in DMSO-treated MV4;11 cells plotted versus the log_2-_HMD (H3K79me2) for +1000 bp from the TSS. Colors signify: red, MLL-AF4 targets^20^; purple, MLL-AF4 targets downregulated by 10 nM pinometostat. **B.** (top) Quantitative measurement of H3K79me2 modification density from ICeChIP-seq of MV4;11 cells treated with 10 nM pinometostat for 7 days contoured over the promoters (−2000 to 2000 bp from the TSS) of indicated gene sets, including genes up-or down-regulated by 10 nM pinometostat, the most highly-expressed genes, MLL-AF4 target genes^20^ as well as those MLL-AF4 targets downregulated by 10 nM pinometostat. (bottom) Heatmaps depicting H3K79me2 density (HMD) for the gene promoter regions shown above ranked by HMD. **C.** Scatterplot of genes in MV4;11 cells downregulated by 10 nM pinometostat depicting log_2_-fold change H3K79me2 HMD (+1000 bp from TSS) versus the log_2_-fold change of the mean normalized FPKM (three independent replicates) for 10 nM pinometostat or DMSO treated cells. Colors signify: red, MLL-AF4 targets^20^; orange, FLT3-ITD upregulated genes^61^; blue, FLT3-ITD downregulated genes^61^; yellow, labeled genes in gray font. **D.** Same as A. but includes 1 nM pinometostat treatment and the subset where this complex spreads^20^. **E.** The FLT3 locus as representative of an MLL-AF4 target^20,22^ downregulated by 10 nM pinometostat, displaying MV4;11 ICeChIP-seq tracks for H3K79me2 10 nM pinometostat 4 and 7 day treatment and H3K27me3 and H3K4me3 tracks from 10 nM pinometostat 7 day treatment as well as DMSO control-treated cells and an RNA-seq track (FPKM) from a single replicate of 10 nM pinometostat 7 day treatment and DMSO-treated cells.

In all gene categories we examined, 10 nM pinometostat dramatically reduced apparent H3K79me2 density in gene bodies, eliminating the sharp peaks near the TSS and proportionally reducing methylation as it tapers toward the 3’ end of the gene body (Figure 3B). The upregulated gene set displayed lower-than-average density both before and after treatment, consistent with the transcriptional upregulation occurring as an indirect effect of the dosing. Whereas the 10 nM pinometostat downregulated genes, 250 highest expressed genes and MLL-AF4 targets all experienced similar reductions in H3K79me2 HMD. The similar reductions in methylation at gene groups that had such different overall responses to gene expression from pinometostat treatment suggests that the expression of some genes is more dependent on H3K79me2-mediated transcriptional activation. Given the modest correlation between H3K79me2 early in the gene body and transcriptional output, we observed an unexpectedly poor linear correlation between fold-change in H3K79me2 HMD versus fold-change in gene expression of differentially expressed genes (R^2^ = 0.13) (Figure S3B). However, comparing the absolute differences in HMD to fold-change of gene expression more clearly reveals some interesting trends (Figure 3C). Those genes with the largest reductions in HMD (including MLL-AF4 targets) are nearly uniformly downregulated though not in proportion to HMD loss. Conversely, MLL-AF4 targets with smaller HMD reductions are more evenly distributed between both up- and downregulated genes. FLT3-ITD-upregulated genes identified in patient samples^61^ have only small reductions in HMD, suggesting their downregulation is not a direct result of HMD loss but, instead, a secondary effect of FLT3 downregulation.

Interestingly, the MLL-AF4 targets downregulated by low-dose pinometostat (Figure 2B) had the largest reductions in H3K79me2 of any gene category examined (Figure 3A). These data show that a subset of MLL-AF4 targets have higher levels of H3K79me2 and greater reductions from DOT1L inhibition and are more dependent on this methylation for even moderate levels of expression. Gene expression sensitivity to low-dose DOT1L inhibition may more accurately define “true” MLL-AF4 target genes whose expression is upregulated by the fusion protein and H3K79me2 hypermethylation than those genes that merely align with MLL1 and AF4 ChIP-seq peaks.

To further define the H3K79me2 depletion trajectory, we also examined the distribution of this modification within gene bodies at an earlier timepoint of pinometostat treatment. Treating MV4;11 cells with 10 nM pinometostat for 4 days had little effect on H3K79me2 HMD at the most highly expressed genes, which likely depend more on DOT1L recruitment by the transcriptional apparatus than by the MLL-fusion protein (Figure 3D). Pinometostat treatment for 4 days diminished the 5’ H3K79me2 peak at genes downregulated by 7-day pinometostat treatment and at MLL-AF4 targets while only slightly reducing H3K79me2 levels within gene bodies of MLL-AF4 targets. Within the gene bodies of 10 nM (7 day) pinometostat-downregulated genes there was actually an increase in H3K79me2 HMD at the 4-day timepoint. This 3’ shift in methylation density away from the transcription start site was even more evident in “MLL-spreading” genes, which showed little reduction in peak methylation levels seen in other groups. The shifting and near total depletion of H3K79me2 density from 4-day and 7-day 10 nM pinometostat treatment respectively, is exemplified by several MLL-AF4 target loci (Figure S6D, S6F, and S6G).

The absence of a correlation between H3K79me2 loss and reductions in gene expression suggests that this modification does not have a universal and proportionate effect on gene activation. Rather, it appears some MLL-AF4 targets have higher levels of H3K79me2 and are more sensitive to its depletion. It is possible that the higher methylation levels result in greater dependence on this modification for gene expression at a subset of MLL-AF4 targets. Given the correlation of H3K79me2 depletion with *FLT3-ITD* expression (Figure 3E), we next sought to determine if these consequences were direct, and whether the functional consequences of DOT1L inhibition can be explained by this pathway.

### MLL-r cells with FLT3-ITD mutations are hypersensitive to both DOT1L and FLT3 inhibition

As our mechanistic analyses relied on MV4;11 cells (*MLL-AF4*, *FLT3^ITD/ITD^)*, we investigated the effects of low dose DOT1L inhibition on 3 other cell lines to determine whether *FLT3-ITD* could account for increased sensitivity to H3K79me2 ablation. Unlike MV4;11, the MOLM13 cell line harbors an *MLL-AF9* translocation and is heterozygous for the *FLT3-ITD* mutation^66^, lesions that have been shown to cooperate to reduce the latency of leukemia onset in mice^23^. We also examined two MLL-translocation cell lines without *FLT3* mutations: THP-1 (*MLL-AF9*); and SEM (*MLL-AF4*). We note that previous studies of DOT1L inhibitor dosing sensitivity of some MLL-r cell lines^26^ could be explained by the *FLT3* mutational status, although given the many other genetic background differences in outgrown cell lines it is reasonable that this correlation was not noted.

We treated all four cell lines with 10 nM pinometostat for 7 days. When comparing each cell line to its counterpart with the same MLL-translocation, those with the *FLT3-ITD* mutation were significantly more sensitive to DOT1L inhibition than those with normal *FLT3* alleles (Figure 4A, left). After 7 days of 10 nM pinometostat treatment MV4;11 viability was drastically reduced by 74% ± 3% while the viability of SEM, its *MLL-AF4* counterpart with intact *FLT3*, was unaltered within experimental error. MOLM13 viability was somewhat reduced (21% ± 3%) while there was no significant difference in the viability of THP-1 cells. As in MV4;11 cells, MOLM13 cells displayed no change in *HOXA9* or *MEIS1* expression under these conditions (Figure S4A).

**Figure 4.**
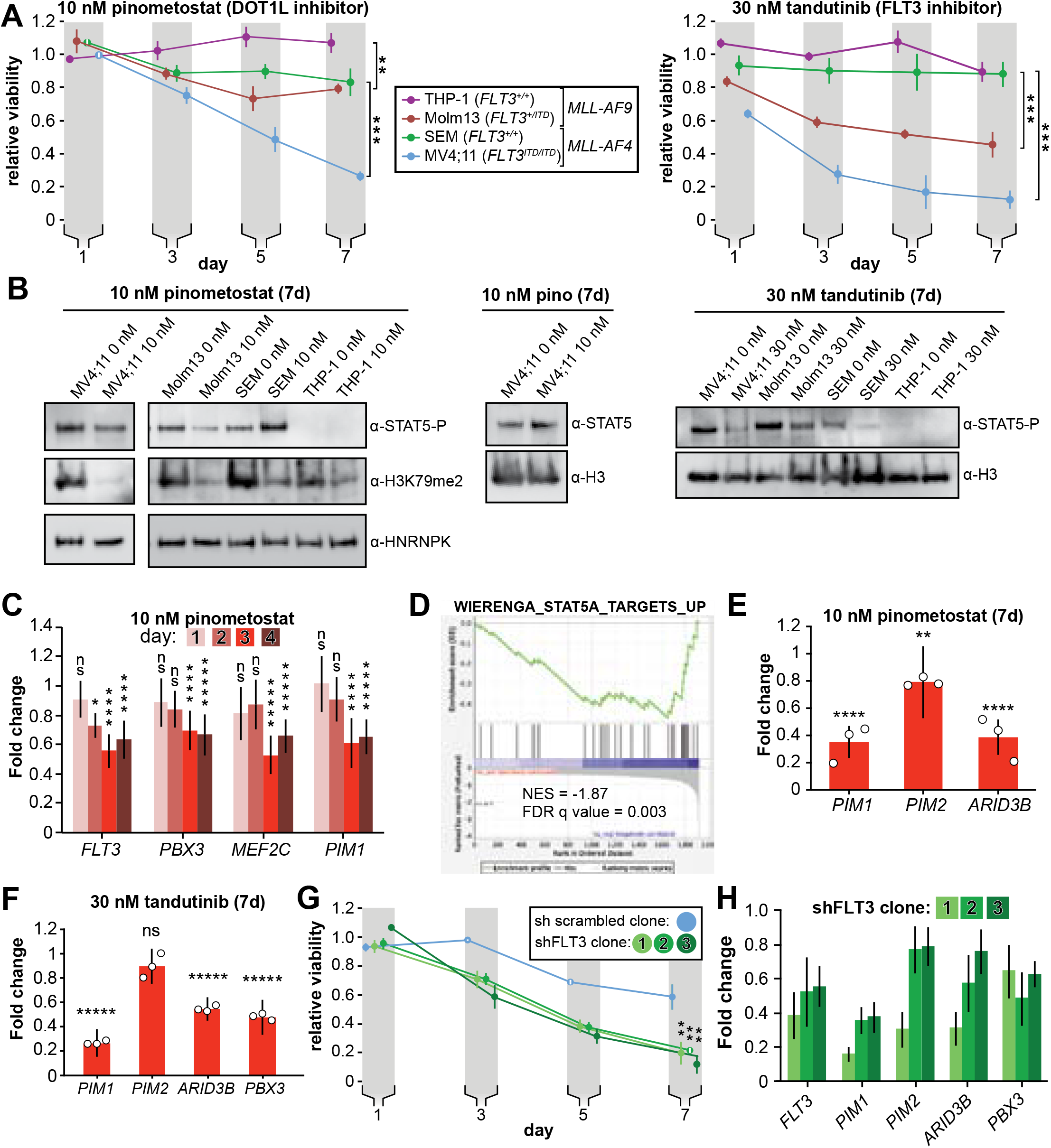
DOT1L inhibition reduces STAT5A activation and downregulates STAT5A targets in *FLT3-ITD* leukemia lines. **A.** MLL-rearranged leukemia lines with genotypes indicated were treated with 10 nM pinometostat (left panel, DOT1L inhibitor) or 30 nM tandutinib (right panel, FLT3 inhibitor MLN518), and relative growth monitored by CellTiter Glo 2.0 assay on the indicated days. Relative viability presented is the mean fraction of luminescence of treated versus side-by-side mock treated cultures (same volume of DMSO) for three independent replicates ± S.E.M. Student’s t-test (** *p* ≤ 0.01, *** *p* ≤ 0.001). **B.** Western blots of phosphorylated STAT5 (active) or total STAT5A with H3 or HNRNPK as loading controls across the cell lines from panel A treated as indicated; H3K79me2 is monitored in pinometostat lines to confirm inhibition. **C.** Time course of gene expression by RT-qPCR, presented as mean fold-change of *FLT3, PBX3, PIM1* and *MEF2C* in MV4;11 cells ± 10 nM pinometostat at each time point indicated ± S.E.M.; n = 3; Student’s t-test (ns *p* > 0.05, * *p* ≤ 0.05, **** *p* ≤ 0.0001, ***** *p* < 0.00001). **D.** Gene Set Enrichment Analysis (GSEA)^46,47^ of the set of downregulated genes in MV4;11 cells ± 10 nM pinometostat compared to genes upregulated by exogenous expression of constitutively active STAT5A in the WIERENGA_STAT5A_TARGETS_GROUP1 gene set^71^ from the MSigDB database. **E-F.** DOT1L and FLT3 inhibition downregulate STAT5A targets in *FLT3-ITD*. RT-qPCR expression analysis presented as mean fold-change ± S.E.M. for the indicated transcript in MV4;11 cells treated with indicated inhibitor versus mock-treatment for 7 days. Student’s t-test (** *p* < 0.01, **** *p* < 0.0001, ***** *p* < 0.00001). **G.** Proliferation assay as in panel A, with 3 clonal populations of MV4;11 cells virally transduced, selected, then induced to express shRNA to FLT3^79^ or a scrambled shRNA^117^ control by 1 μg/mL doxycycline. Means of fractional viability relative to uninduced cells ± S.E.M. are shown for 3 independent experiments; Student’s t-test (** *p* < 0.01). **H.** RT-qPCR analysis of *PIM1, PIM2* and *ARID3B* expression in MV4;11 cells expressing an inducible shRNA targeting FLT3^79^ for 7 days. Results are depicted as fold-change expression of control cells expressing shRNA to GFP^118^.

If the heightened sensitivity of MLL-r cell lines to DOT1L inhibition is indeed mediated by reduced *FLT3-ITD* expression, then we would expect to see a similar heightened sensitivity to disruption of FLT3 signaling. The small molecule tandutinib (MLN518) inhibits FLT3 kinase activity, severely reducing phosphorylation-mediated activation of downstream targets such as STAT5A^67^. We treated our MLL-r cell lines with 30 nM tandutinib for 7 days. As with the DOT1L inhibition experiments, cell lines with *FLT3-ITD* mutations were significantly more susceptible to the inhibitor’s effects (Figure 4A, right). Given the variety of other genetic differences amongst these cell lines, these observations can at best be taken as consistent with the hypothesis that the co-occurring *FLT3-ITD* mutations may sensitize MLL-r leukemias to DOT1L inhibition, motivating us to seek more direct examination of FLT3 signaling.

### Impaired FLT3 signaling by DOT1L inhibition culminates in reduced transcription of STAT5A target genes

The *FLT3-ITD* mutation allows FLT3 to phosphorylate STAT5A, a transcription factor that is not activated by wild type FLT3^68^. This aberrant STAT5A phosphorylation licenses translocation to the nucleus to drive target gene transcription, resulting in a hyperproliferative state necessary for leukemia cell survival^69,70^. We hypothesized that *FLT3-ITD* downregulation by DOT1L inhibition would thereby reduce STAT5A phosphorylation. Indeed, pinometostat treatment reduced STAT5 phosphorylation in MV4;11 cells without affecting STAT5A protein levels (Figure 4B). Pinometostat treatment slightly reduced STAT5 phosphorylation in MOLM13 cells, consistent with the lower *FLT3-ITD* allele dose, whereas lines with wild type *FLT3* (THP-1, SEM) did not display these effects. As a point of direct comparison, small molecule inhibition of FLT3 signaling yielded markedly reduced STAT5 phosphorylation in lines bearing the *FLT3-ITD* (MV4;11 and MOLM13), with a more modest reduction in SEM cells while phospho-STAT5 was barely detectable in THP-1 cells (Figure 4B).

To examine whether FLT3 effects precede other pro-proliferation pathways, we obtained more granular expression kinetics of several downregulated MLL-AF4 targets that have been implicated in leukemogenesis. Expression of *FLT3, PBX3, PIM1* and *MEF2C* was significantly reduced after 72 hours treatment with pinometostat (Figure 4C), however, *FLT3* was the only gene whose expression was reduced 48 h after treatment, suggesting it is more sensitive to H3K79me2 reductions than the others examined. Though *FLT3* and *MEF2C* are targets of the HOXA9-MEIS1-PBX3 complex, these genes are all targets of the MLL-fusion protein^20^. The reduction in *FLT3* expression in advance of decreased *PBX3* or *MEF2C* expression lends tentative support to the possibility that DOT1L inhibition directly affects *FLT3* gene expression independently of *PBX3* or *MEF2C*.

Given the early reductions in FLT3-ITD expression and reduced phosphorylation of its target STAT5A, we hypothesized that the pinometostat-induced reductions in proliferation were due to a loss of STAT5A signaling. We performed GSEA^46,47^ with the pinometostat-downregulated genes and genes upregulated by STAT5A overexpression in human CD34+ hematopoietic progenitors^71^ and observed a negative correlation indicative of significant pathway overlap (NES = −1.87, FDR = 0.003, Figure 4D). We then reexamined our RNA-seq data for previously described STAT5A target genes downregulated by pinometostat and found several, including *PIM1* and *ARID3B*^72,73^(Figure S4C). The PIM proteins are a family of 3 protooncogene serine/tyrosine kinases (PIM1-3) that are upregulated in, and indicative of poor prognosis in leukemia, prostate, mesothelioma and other cancers^55,72,74–77^. However, only *PIM1* and *PIM2* expression is increased in FLT3 inhibitor-resistant *FLT3-ITD* patient samples and exogenous expression of either *PIM1* or *PIM2* can rescue proliferation defects caused by loss of FLT3 activity in MOLM14 cells (*MLL-AF9*, *FLT3-ITD* heterozygous)^78,79^. Although *PIM1* and *PIM2* are both downregulated in our RNA-seq analysis (Figure S4C) we observed a much greater reduction in *PIM1* expression by RT-qPCR (Figure 4E). Similarly, treating MV4;11 cells with tandutinib (FLT3 inhibitor) resulted in downregulation of *PIM1*, *ARID3B* and *PBX3* but not *PIM2* (Figure 4F). Treating MOLM13 cells with pinometostat also reduced expression of *MEF2C, FLT3* and *PIM1,* but caused no change in *PBX3* expression (Figure S4D).

If the FLT3 and DOT1L inhibitors have overlapping functions through inhibition or downregulation of *FLT3,* respectively, then we could potentially observe synergy in the effects on MV4;11 proliferation if we treated with both inhibitors simultaneously. The DOT1L inhibitor has a delayed effect compared to the PIM1 and FLT3 inhibitors, which complicates comparisons, but nonetheless, we observed small but significant differences in proliferation when using inhibitors singly or in combination at day 7 (Figures S4E and S4F).

To directly interrogate the effects of *FLT3* on MLL-r leukemia proliferation without complications from different genetic backgrounds, we used viral transduction to insert a tet-inducible shRNA targeting *FLT3* into MV4;11 cells. With modest knockdown of *FLT3* (Figure 4H) we observed significant reductions in the proliferation of 3 different clonal lines as compared to a scrambled shRNA (Figure 4G). *FLT3* knockdown reduces MV4;11 proliferation and STAT5A phosphorylation (Figure S4G), analogous to the effects of pinometostat treatment. Akin to the DOT1L and FLT3 inhibitors (Figure 4E-F), *FLT3* knockdown also significantly reduced the expression of the STAT5A target genes *PIM1* and *ARID3B*, with *PIM2* expression reduced in only 1 of 3 clones (Figure 4H). Interestingly, *FLT3* knockdown also resulted in *PBX3* downregulation, suggesting that FLT3 can regulate the expression of this oncogenic transcription factor, in line with previous observations^61^. Unfortunately, overexpression of *FLT3-ITD* for an attempted rescue of DOT1L inhibition proved technically challenging, as retrovirally introduced ectopic expression was rapidly silenced or dropped out during selection as has been observed in other contexts^44^. To further interrogate this pathway’s functional significance, we sought to perturb signaling downstream of FLT3-ITD via STAT5A alterations.

### Overexpression of constitutively active STAT5A rescues proliferation and reductions in gene expression caused by DOT1L inhibition

To potentiate STAT5A activity, we overexpressed a constitutively active STAT5A mutant to examine whether this could counteract the reduction of upstream FLT3-ITD levels by DOT1L inhibition. STAT5A is “activated” through phosphorylation at multiple sites, facilitating translocation into the nucleus and activation of gene targets. Previous work showed that H299R and S711F mutations create a constitutively active murine *Stat5a* able to activate target genes independently of upstream signaling^69^, which phenocopies the effects of exogenous *FLT3-ITD* expression including hyperproliferation and inhibition of myeloid maturation^80^. We used a lentiviral system to generate individual MV4;11 clonal cell lines with stably-incorporated, inducible human *STAT5A* mutated at the corresponding residues H298R and S710F (*STAT5A-CA*), all of which exhibit several-fold induction with doxycycline (Figure 5A and Figure S5A). Ectopic expression of *STAT5A-CA* was able to partially rescue proliferation when challenged with 30 nM FLT3 inhibitor tandutinib, confirming the capacity of this mutant to complement impaired FLT3-ITD signaling (Figure S5B).

**Figure 5.**
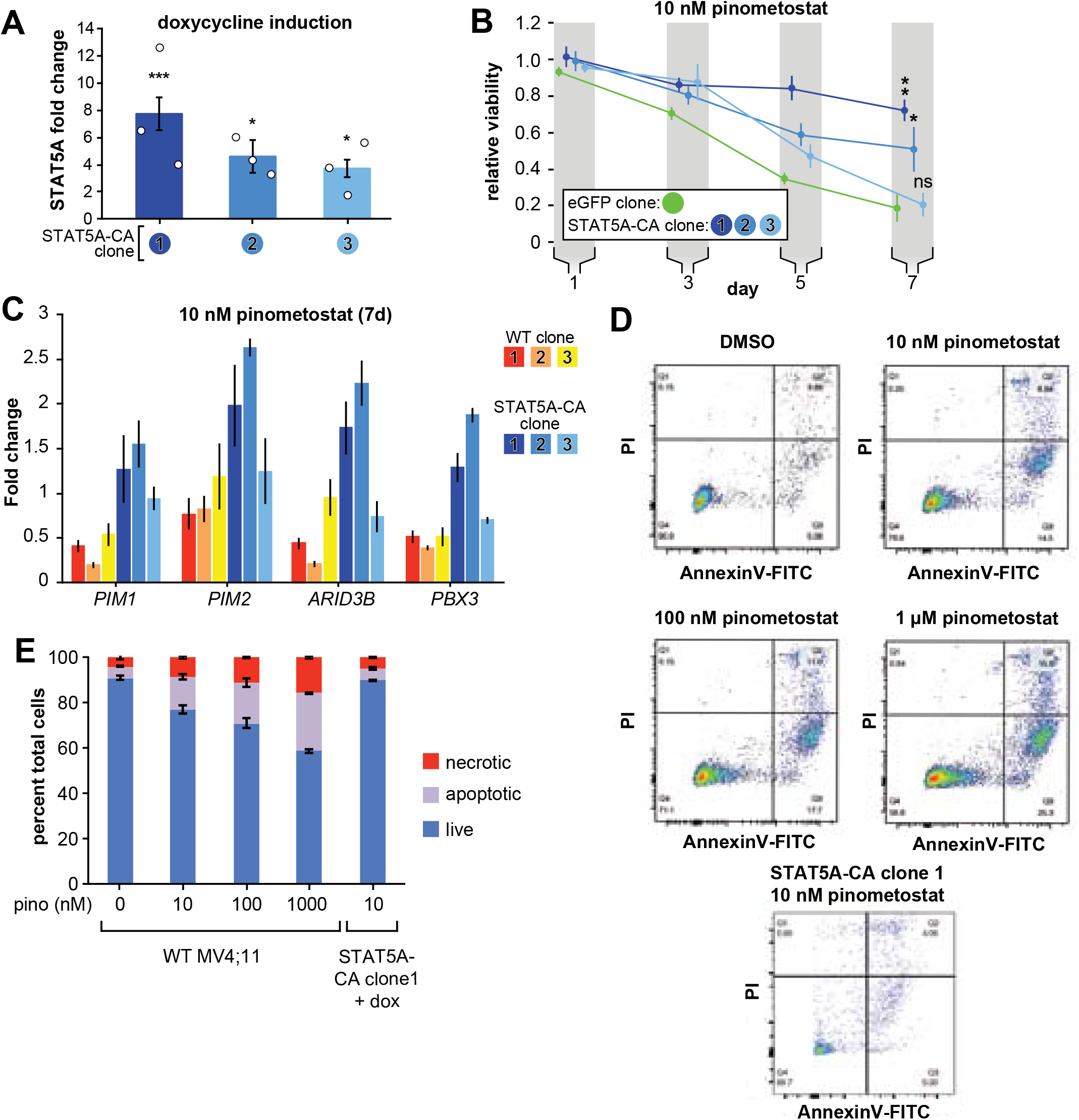
Exogenous expression of constitutively active *STAT5A* partially rescues proliferation and gene expression effects of DOT1L inhibition. **A.** RT-qPCR analysis of *STAT5A* expression from 3 monoclonal isolates of MV4;11 cells virally transduced with a tet-inducible constitutively active *STAT5A* (*STAT5A-CA*) depicted as fold-change over untransduced cells with standard error of the mean. Student’s t-test (* *p* < 0.05, *** *p* < 0.001). **B.** Proliferation assay of MV4;11 clonal isolates from panel A. induced to express *STAT5A-CA* or eGFP with 1 μg/mL doxycycline and treated concomitantly with 10 nM pinometostat. We determined the fractional viability of each clone as the luminescence from a CellTiter Glo 2.0 assay with pinometostat-treatment normalized to DMSO-treated cells, both induced to express *STAT5A-CA or eGFP*, to accommodate for any potential increases in viability. Means ± SE are shown for 3 independent experiments with Student’s t-test for significance of day 7 values (**** *p* ≤ 0.0001). **C.** Gene expression analysis by RT-qPCR of STAT5A target genes in WT MV4;11 cells or MV4;11 *STAT5A-CA* clones from A. induced with 1 μg/mL doxycycline and treated with 10 nM pinometostat for 7 days. Results are displayed as fold-change over DMSO-treated WT cells. **D.** Quantitative measurement by flow cytometry of live, apoptotic (Annexin V-FITC) and necrotic cells (propidium iodide) of WT MV4;11 cells or cells exogenously expressing *STAT5A-CA* (clone 1) and treated with increasing concentrations of pinometostat. Images of gated FITC vs. PI signal are shown for 1 of 3 independent experiments that are quantified in the bar plot in **E**.

Remarkably, *STAT5A-CA* overexpression also rescued pinometostat-induced proliferation reductions (Figure 5B) in proportion to each clone’s *STAT5A-CA* expression level (Figure 5A). Clone 3 was unable to rescue proliferation substantially, possibly because it had the lowest expression of *STAT5A/STAT5A-CA* (Figure 5A). As another control, we similarly overexpressed MEF2C, yet it displayed no effect on the viability of MV4;11 cells treated with 10 nM pinometostat (Figure S5C).

To gain a molecular understanding of how ectopic *STAT5A-CA* expression could rescue proliferation of inhibitor-treated cells, we measured expression of the STAT5A targets *PIM1*, *PIM2* and *ARID3B* by RT-qPCR. Expression of *STAT5A-CA* restored expression of *PIM1*, *PIM2* and *ARID3B* in both DOT1L inhibitor- and FLT3 inhibitor-treated MV4;11 cells (Figure 5C and S5D).

Because ectopic expression of *STAT5A-CA* is able to rescue proliferation of MV4;11 cells and the expression of STAT5A targets including the anti-apoptotic *PIM1* oncogene, we examined whether *STAT5A-CA* overexpression could rescue MV4;11 cells from apoptosis. A previous study observed that ~30% of MV4;11 cells treated with 1 μM pinometostat for 6 days were undergoing apoptosis^26^. We analyzed apoptosis in MV4;11 cells treated with increasing concentrations of pinometostat for 7 days (Figures 5D and E). We observed 25.5% ± 0.3% apoptotic cells when treating with 1 μM pinometostat and a still sizeable proportion (15% ± 1%) of apoptotic cells when treating with just 10 nM pinometostat. Yet upon treatment of STAT5A-CA clone 1 with 10 nM pinometostat for 7 days, we observed no significant induction of apoptosis as compared to the DMSO control (Figure 5D and E). Thus, we concluded that recovering STAT5A function can rescue MV4;11 cells from apoptosis induced by 10 nM pinometostat. It is striking that despite marked gene expression changes caused by low-dose DOT1L inhibition, one signaling pathway, FLT3-ITD to STAT5A, is able to account for the bulk of the phenotypic and molecular changes we measured. Given that the rescue was nevertheless incomplete, we investigated other potential secondary contributors to the proliferation and gene expression consequences of low-dose DOT1L inhibition.

### An ancillary DOT1L-dependent pathway limits proliferation through PRC2 signaling

Although H3K79me2 potentiates transcription, our RNA-seq analysis revealed the upregulation of thousands of genes when treating with pinometostat. One potential explanation for this effect is the downregulation of the repressive PRC2 complex members *EZH2* and *EED* and consequent reductions in global levels of the transcriptionally repressive H3K27me3 mark (Figures 6A-B, S6A). PRC2 deposits the facultative heterochromatin H3K27me3 modification and, though antagonistic to MLL1 and H3K4me3 deposition^81^, is necessary for MLL-r leukemogenesis^82–84^.

**Figure 6.**
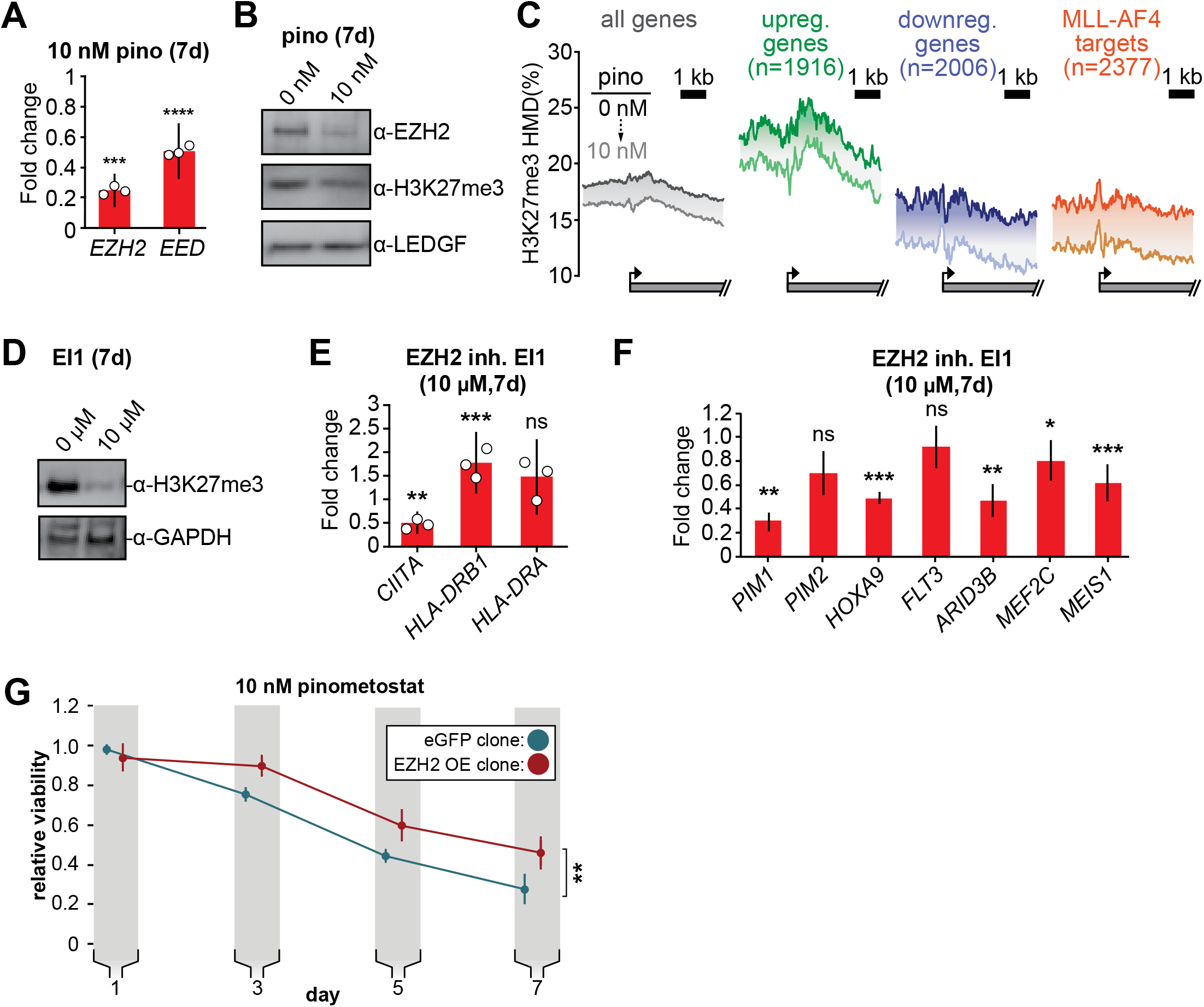
PRC2 function is an ancillary pathway dependent on DOT1L and necessary for leukemia proliferation. **A.** RT-qPCR analysis of the components of the polycomb complex *EZH2* and *EED* expression in MV4;11 cells ± 10 nM pinometostat for 7 days. Results are displayed as mean fold-change vs. DMSO-treated cells ± S.E.M. of three independent experiments. Student’s t-test for significance (*** *p* < 0.001, **** *p* < 0.0001). **B.** Western blot of EZH2, H3K27me3 and LEDGF as loading control in MV4;11 cells treated ± 10 nM pinometostat for 7 days. **C.** Quantitative Ice-ChIP-seq from MV4;11 cells treated with 10 nM pinometostat for 7 days displaying H3K27me3 histone methylation density contoured over promoters from −2000 to +4000 of the TSS of either all expressed genes, genes up-or downregulated by 10 nM pinometostat or MLL-AF4 target genes^20^. **D.** Western blot for H3K27me3 with GAPDH as a loading control in MV4;11 cells treated with EI1 for 7 days. **E.** RT-qPCR analysis of MHC class II genes and master regulator *CIITA* expression from MV4;11 cells ± 10 μM EZH2 inhibitor EI1. Results are displayed as mean fold-change vs. DMSO-treated cells ± S.E.M. of three independent experiments. Student’s t-test for significance (** *p* = 0.01, *** *p* = 0.001). **F.** Fold change of RT-qPCR analysis of gene expression MV4;11 cells ± 10 μM EZH2 inhibitor EI1. Results are the average 3 independent experiments ± S.E.M. Student’s t-test (* *p* < 0.05, ** *p* < 0.01, *** *p* = 0.001). **G.** Proliferation assay of MV4;11 cells virally transduced with tet-inducible EZH2 or eGFP treated with 10 nM pinometostat and induced with 1 μg/mL doxycycline to express EZH2 or eGFP for 7 days showing the luminescence fraction of inhibited over uninhibited from a CellTiter Glo 2.0 assay. Means ± SE are shown for 3 independent experiments. Student’s t-test of day 7 values (** *p* = 0.01).

Analysis by quantitative ICeChIP revealed that 10 nM pinometostat decreased H3K27me3 genome-wide (Figure 6C). Promoter H3K27me3 levels are reduced by 2-5% on average with more pronounced decreases observed among downregulated genes and MLL-AF4 targets than upregulated or all genes (Figure 6C). However, H3K27me3 levels in untreated cells were much higher in pinometostat-upregulated genes, perhaps indicating that these genes are more reliant on PRC2 to buffer their expression. H3K27me3 levels are lower throughout gene bodies in DOT1L inhibited cells, as apparent at individual loci (Figure 3E, S6D-G).

We next sought to interrogate the functional impact of the PRC2 signaling axis by experimental perturbation. As PRC2 is necessary for repression of *IFNG* (IFN-γ) and proper differentiation in T-cells^88^, we wondered if the upregulated genes found in our RNA-seq analysis, many of which are components of the IFN-γ-response, were upregulated as a result of a loss of H3K27me3-mediated repression. To investigate this possibility, we treated MV4;11 cells with 10 μM EI1 EZH2 inhibitor ^87^ and observed dramatically reduced global H3K27me3 (Figure 6D) and proliferation (Figure S6B), consistent with previously observed sensitivities of MOLM13 and MV4;11^85^. EI1 treatment had comparatively little effect on the class of genes massively overexpressed during DOT1L inhibition (Figure 6E, compare to Figures 2D and E). Surprisingly, EZH2 inhibition downregulated *HOXA9* and *MEIS1* expression (which only occurs with higher doses of pinometostat^26^), with no changes in *FLT3* expression (Figure 6F) or STAT5 phosphorylation (Figure S6C). The greater reduction in global H3K27me3 from 10 μM EI1 than 10 nM pinometostat may account for the lack of effect on *HOXA9* and *MEIS1* expression by pinometostat. Collectively, these data argue that the PRC2 pathway is largely independent of the FLT3-ITD-STAT5A pathway, culminating in distinct target gene expression consequences, that may converge for only a few targets, such as *PIM1* and *ARID3B*.

Next, we queried the functional consequences of rescuing EZH2 expression in the context of low-dose DOT1L inhibition. Inducible overexpression of *EZH2* was only able to partially rescue proliferation in M4;11 cells treated with pinometostat, suggesting that a small portion of the effects on MV4;11 viability is due to reduced PRC2 function (Figure 6G). The nearly complete rescue from intervening in the FLT3-ITD-STAT5A pathway compared to the modest rescue from PRC2, suggests that the former is the predominant source of pinometostat-induced effects on proliferation in this leukemia background.

### STAT5A-CA overexpression rescues the viability of MV4;11 cells treated with MLL1 inhibitors

Our observations suggest that most of the toxicity from low-dose DOT1L inhibition in MLL-r, *FLT3-ITD+* leukemia cell lines stems from downregulation of *FLT3* and subsequent loss of STAT5A phosphorylation. We wanted to know if this effect was specific to H3K79me2 depletion, or attributable to disruption of MLL-fusion-induced gene activation. To distinguish between these two mechanisms, we employed small-molecule MLL1 inhibitors, potent and effective treatments for MLL-r leukemia^15,86^, as orthologous means of disrupting MLL-fusion function. These compounds inhibit MLL1 in different ways but both disrupt the leukemic gene expression profile, specifically downregulating the oncogenes *HOXA9*, *MEIS1*, *FLT3* and *BCL2*^15,86^. MI-503 competitively antagonizes binding of MENIN to MLL1, an interaction that is necessary for MLL-fusion complex localization to target genes and leukemogenesis^21,92,^. Another small molecule, MM-401 inhibits the methyltransferase activity of MLL1 by disrupting its interaction with WDR5, a complex member necessary for full enzymatic activity of MLL1 but not MLL2-4 or SET1 complexes^15^. We treated MLL-r cell lines with low concentrations of MI-503 or MM-401 and observed greater reductions in the proliferation of *MLL-r*, *FLT3-ITD+* cells than their WT *FLT3* counterparts (Figure 7A and B).

**Figure 7.**
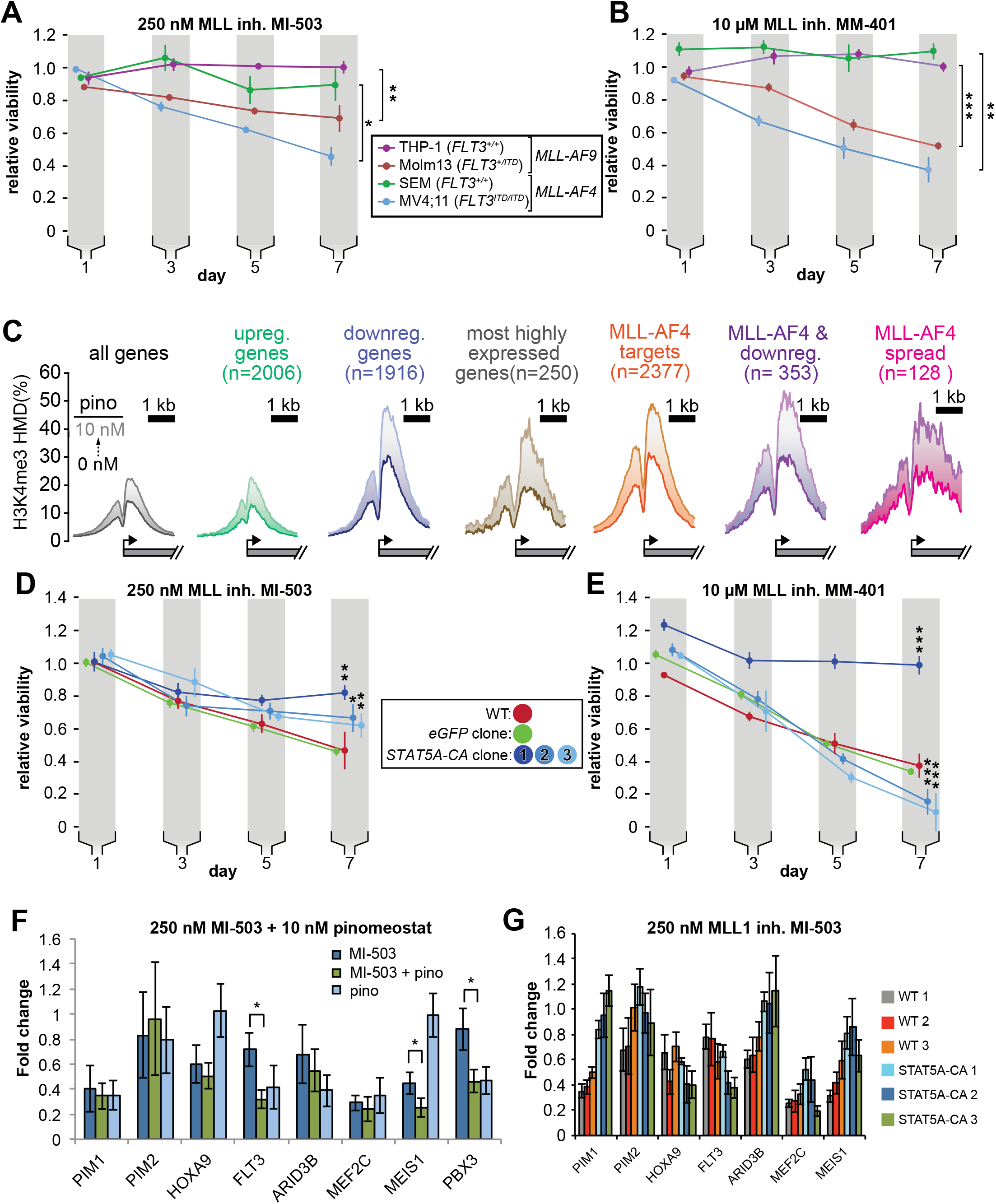
STAT5A-CA overexpression rescues the viability of MV4;11 cells treated with MLL1 inhibitors. Proliferation assay of MLL-r cell lines treated with **A.** 250 nM MI-503 (MLL1-Menin interaction inhibitor) or **B.** 10 μM MM-401 (MLL1 histone methyltransferase inhibitor) for 7 days. Viability was measured by CellTiter Glo 2.0 assay and results are displayed as the fraction of luminescence of inhibitor-treated over DMSO-treated cells. Means ± SE are shown for 3 independent experiments. Student’s t-test (* *p* < 0.05, ** *p* < 0.01, *** *p* < 0.001). **C.** H3K4me3 histone methylation density from −2000 to +2000 of the TSS from quantitative ICeChIP-seq from MV4;11 cells treated with 10 nM pinometostat for 7 days for genes up-or down-regulated by 10 nM pinometostat, the most highly-expressed genes, MLL-AF4 target genes^20^ as well as those MLL-AF4 targets downregulated by 10 nM pinometostat. **D. and E.** Proliferation assay of MV4;11 *STAT5A-CA* clonal isolates induced to express *STAT5A-CA* or eGFP with 1 μg/mL doxycycline and treated with **D.** 250 nM MI-503 or **E.** 10 μM MM-401. Viability was measured and results displayed as in A and B. Means ± SE are shown for 3 independent experiments. Student’s t-test (* *p* < 0.05, ** *p* < 0.01, *** *p* < 0.001). **F.** Gene expression analysis by RT-qPCR of MLL-fusion and STAT5a targets in MV4;11 cells treated with 250 nM MI-503 MLL1 inhibitor, 10 nM pinometostat DOT1L inhibitor or a combination for 7 days. Means ± S.E.M. are shown for 3 independent experiments (* *p* < 0.05). **G.** Gene expression analysis by RT-qPCR of MLL-fusion and STAT5A targets in WT and STAT5A-CA MV4;11 cells treated with 250 nM MI-503 MLL1 inhibitor for 7 days. Means ± S.E.M. are shown for technical replicates of individual experiments.

Given the similar effects of DOT1L and MLL1 inhibitors on MLL-r cell proliferation and gene expression, that both histone modifications are involved in transcriptional activation and the extensive literature describing dynamic cross-talk between chromatin modifications^81,89–91^ we were curious as to how perturbations in H3K79 methylation might affect the distribution of H3K4me3. In order to accurately quantify histone methylation and observe differences in modification densities we performed ICeChIP-seq for H3K4me3 in MV4;11 cells treated with pinometostat. H3K4me3 is deposited at promoters during active transcriptional initiation and promotes gene expression through several established mechanisms^92–94^. Surprisingly, pinometostat treatment increased H3K4me3 at transcription start sites (TSS’s) genome-wide, with the largest increases at genes downregulated by pinometostat (Figure 7C). Pinometostat-downregulated MLL-AF4 targets had the highest H3K4me3 levels of all gene categories examined (Figure 7C), not only at the TSS but spreading downstream into the gene body, suggesting that the MLL-fusion protein is driving this increase. Potentially indicative of a previously undescribed form of antagonistic histone mark cross-talk, we observe a striking anti-correlation between pinometostat-induced reductions in H3K79me2 and increases in H3K4me3, which was most evident at MLL-AF4 targets downregulated by 10 nM pinomeostat. Despite gains of the H3K4me3 mark during treatment, these genes are downregulated, consistent with a decoupling of active transcription initiation from productive elongation, the latter of which is more effectively correlated with H3K79me2 and H3K36me3^65^.

Intriguingly, the putative antagonism between modifications is not apparent in global H3K4me3 levels during DOT1L inhibition (Figure S7A). However, reductions in H3K4me3 from MLL1 inhibitor treatment are also not readily apparent by Western blot, similar to what has been observed in other studies^15^ (Figure S7B). Conversely, global increases in H3K79me2 are more pronounced when treating cells with the MLL1 inhibitors (Figure S7C). Treatment with the MLL1 inhibitors also reduced STAT5A phosphorylation, suggesting that this orthologous means of disrupting MLL-fusion gene activation also reduces FLT3-ITD/STAT5A signaling (Figure S7C).

As with the DOT1L and FLT3 inhibitors, overexpression of *STAT5A-CA* was able to partially rescue survival of MV4;11 cells treated with MI-503 (Figure 7D), with the degree of rescue corresponding to the amount of *STAT5A* expression in each clone (Figure 5A). When treated with the MM-401 inhibitor, STAT5A-CA clone 1 (with the highest exogenous *STAT5A-CA* expression), completely rescued proliferation (Figure 7E). Unexpectedly, clones 2 and 3, that express *STAT5A-CA* at lower levels both displayed reduced proliferation when treated with MM-401 compared to WT or GFP expressing cells (Figure 7E).

We observed an additive effect when MV4;11 cells were co-treated with the MLL1 and DOT1L inhibitors (Figure S7D), suggesting that the inhibitors affect different sets of genes through different mechanisms, or have an additive effect on the same genes. To distinguish between these two models, we compared gene expression of several MLL-AF4 and STAT5A targets in MV4;11 cells treated with MI-503 alone or MI-503 with pinometostat for 7 days (Figure 7F). Akin to low dose DOT1L inhibitor treatment, MI-503 reduced expression of *FLT3*, *MEF2C, ARID3B* and *PIM1*. The reduction in *FLT3* expression was only 30% ± 10% but doubled to 60% ± 10% when both inhibitors were used, recapitulating the 60% reduction observed with pinometostat alone. MI-503 had no significant effect on *PBX3* expression but both inhibitors reduced *PBX3* expression to 50% ± 10%, the same as the DOT1L inhibitor alone.

However, unlike low dose pinometostat, MI-503 treatment starkly reduced expression of *HOXA9* and *MEIS1* and combination treatment further reduced *MEIS1* expression from 40% to 30%. In summation, low dose MLL1 and DOT1L inhibitors downregulate different, yet partially overlapping sets of genes (with *FLT3*, *MEF2C*, and *PIM1* in common), that are necessary for MLL-rearranged leukemia.

We wondered whether the *STAT5A-CA*-mediated rescue of proliferation in MV4;11 cells treated with MI-503 coincided with a rescue of STAT5A target genes. We examined expression of these targets in our 3 *STAT5A-CA* clones after treating cells with MI-503 for 7 days and observed increased the expression of the STAT5A target genes *PIM1, PIM2* and *ARID3B* (Figure 7G). Collectively, these data suggest that downregulation of *FLT3-ITD*, and crucially, reductions in STAT5A phosphorylation and gene activation are more sensitive to perturbations of MLL-fusion-mediated gene activation and are the main source of inhibitor effects on leukemia cell survival when expression of the canonical MLL-r proliferation mediators *HOXA9* and *MEIS1* are not substantially affected (model, Figure S7E).

## Discussion

Little is known about why MLL-r leukemia cell lines have such disparate sensitivities to DOT1L inhibitors or how MLL-fusions might cooperate with co-occurring lesions. By investigating the effects of a DOT1L inhibitor at a low, as yet unexplored concentration, we revealed that MLL-r cell lines carrying *FLT3-ITD* lesions are more sensitive to DOT1L inhibition. We observed that a subset of MLL-AF4 targets, including *FLT3,* have aberrantly high H3K79me2 and that low-dose inhibitor treatment downregulates these genes, dramatically depleting H3K79me2, while resulting in increased H3K4me3 at promoters and reduced H3K27me3 genome-wide. Our findings illustrate how MLL-fusions can cooperate mechanistically with *FLT3-ITD* mutations to facilitate leukemogenesis and how PRC2 function is necessary for that disease state. FLT3-ITD-mediated STAT5A activation is crucial to the MLL-AF4 expression profile, potentially through direct interaction of STAT5A with HOXA9 and coactivation of some targets such as *PIM1*.

### The FLT3-ITD signaling pathway accounts for the bulk of low-dose DOT1L inhibitor toxicity

A subset of MLL-AF4 targets were downregulated by low-dose DOT1L inhibition and the FLT3 locus was impacted earlier than other MLL-AF4 targets. *FLT3* expression was downregulated after only 2 days of low-dose pinometostat treatment, coinciding with reduced proliferation, increased apoptosis and gene expression changes consistent with differentiation. Reductions in *FLT3-ITD* expression precede reductions in other MLL-AF4 targets including *PBX3* and *MEF2C*, arguing that these effects are more primary or sensitive to DOT1L function. Although PBX3 interacts with both HOXA9 and MEIS1 to facilitate leukemogenesis and regulate the expression of common targets including *FLT3*^53,58^ we observed that *PBX3* expression could also be reduced by either FLT3 knockdown or inhibition (Figure 4F and H). These results are in agreement with previous findings that *PBX3* was significantly upregulated in *FLT3-ITD^+^* compared to WT *FLT3,* karyotypically normal AML patient samples^61^.

The FLT3 receptor has an outsized effect on myeloid differentiation and proliferation through its regulation of several myeloid transcription factors^55,95^, accounting for its predominance in AML patients^51,54,55^. Although stable transfection of *FLT3-ITD* has been observed to downregulate the PU.1 and C/EBPα transcription factors and regulators of myeloid differentiation^55^, we detected no discernable change in *SPI1* (PU.1) expression and a surprising ~2-fold downregulation of *CEBPA* (C/EBPα) in MV4;11 cells (Figure S1D) treated with low-dose pinometostat. Much of the *FLT3-ITD*-driven effects on proliferation, inhibition of apoptosis and differentiation have been attributed to the activation of STAT5A^55,80,95–97^. Constitutively active *Stat5a* can render mouse Ba/F3 cells growth factor-independent and resistant to apoptosis through upregulation of the *Pim1-2* protooncogenes^76,110,111^. We observe that 10 nM pinometostat downregulates *FLT3-ITD* with concomitant reductions in STAT5A phosphorylation and diminished expression of the STAT5A target genes *PIM1* and *ARID3B,* suggesting that low-dose DOT1L inhibition is able to disrupt FLT3-ITD-mediated signaling and downstream oncogene activation.

Exogenous expression of constitutively active human *STAT5A* (*STAT5A-CA*) in MV4;11 cells treated with 10 nM pinometostat rescues cells from apoptosis, almost completely rescues proliferation, and restores *PIM1* and *ARID3B* gene expression, suggesting that most of the toxicity from low-dose DOT1L inhibition is through loss of STAT5A activation. The ability of ectopic *STAT5A-CA* expression to rescue orthologous perturbations to MLL-fusion-mediated gene activation and proliferation from MLL1 inhibitors suggests that STAT5A activation is necessary for leukemogenesis and maintenance of the proliferative gene expression profile including *PIM1* in this context. Interestingly, *PIM1* is a downstream target of both FLT3-ITD and HOXA9^72,100^. Though both factors regulate *PIM1* expression, the FLT3-ITD axis is more sensitive and is responsible for *PIM1* downregulation with low-dose DOT1L inhibitor treatment in MLL-r leukemia also bearing the *FLT3-ITD* mutation. FLT3-ITD*-*mediated activation of STAT5A may promote HOXA9 localization to the *PIM1* locus or complement it, thereby facilitating expression of this common target and leukemogenesis.

*PIM1* activation by both STAT5A and HOXA9 represents a common coregulation scenario for these hematopoietic transcription factors. Indeed, De Bock et al. discovered that HOXA9 binding sites have significant overlap with STAT5A, PBX3 and C/EBP targets genome-wide^101^. We observed downregulation of both *PBX3* and *C/EBPA* by 10 nM DOT1L inhibition. It is possible that the dependence of MLL-r, *FLT3-ITD^+^* leukemia on *FLT3-ITD* expression may be due to HOXA9 requiring STAT5A and/or PBX3 and C/EBPA to cooperatively bind select target genes. Huang et al. found that HOXA9 and MEIS1 preferentially localized to enhancer regions enriched with STAT5 binding motifs^100^ and identified STAT5A and C/EBPA in complex with HOXA9. Furthermore, *HOXA9* knockdown reduced STAT5A binding at common target sites^100^. If HOXA9 depends on STAT5A for chromatin localization then low dose DOT1L inhibition may reduce HOXA9 binding at enhancer regions, reducing HOXA9 target gene activation without affecting *HOXA9* expression.

In addition to gene activation, STAT5A phosphorylation also results in gene repression, modulating the immune response and differentiation^80,102^. Viral transduction of constitutively active *Stat5a* affects T cell differentiation by repressing IFN-γ production^102,103^. We found 2007 genes upregulated with 10 nM pinometostat treatment, including many MHC class II genes with large fold-changes that significantly overlapped with a set of genes consistently downregulated in FLT3-ITD+ (KN) leukemia samples^61^. Indeed, GO analysis of the pinometostat-upregulated genes indicated significant enrichment for the “IFN-γ-mediated signaling pathway” and other immune-related categories (Figure S2A). Despite the increase in expression of IFN-γ-regulated genes we saw barely measurable levels of *IFNG* (IFN-γ) and no increase in expression with pinometostat treatment (Figure S2B). Many components of the IFN-γ pathway, such as IRF4 and IRF5 are involved in macrophage differentiation, a functional consequence of *DOT1L* deletion or inhibition that has been observed in other studies^21,33,48,104^. With pinometostat treatment we observed upregulation of *CSF1R* and *CSF3R,* targets of IRF4 and critical signaling inducers of macrophage and neutrophil differentiation, respectively (Figure S2D)^48,49^. Additionally, expression increases in the macrophage cell surface markers *ITGAM* (CD11b), *ITGAX* (CD11c) and *CD86* suggest these cells are differentiating to a more macrophage-like state (Figure S2C), consistent with previous observations from DOT1L deletion and from a study using the DOT1L inhibitor EPZ004777^21,33^.

### Extensive histone modification cross-talk contributes to the survival of MLL-r, FLT3-ITD+ leukemia

*FLT3* is part of a subset of MLL-AF4 targets that are more sensitive to reductions in H3K79me2 than even the *HOXA9* and *MEIS1* oncogenes. We observed that MLL-AF4 targets^20^ that are downregulated by 10 nM pinometostat have higher levels of H3K79me2 than even the most highly expressed genes and show the largest reductions in methylation when treated with pinometostat. (Figure 3A). The greater reductions in H3K79me2 levels at downregulated genes is likely a contributing factor to their loss of gene expression. H3K79me2 hypermethylation antagonizes SIRT1 localization to MLL-AF4 targets, preventing H3K9ac and H3K16ac deacetylation, thereby facilitating gene expression^24^. However, there are stark differences in methylation density and susceptibility to DOT1L inhibition even among MLL-fusion targets. MLL-AF4 “spreading” genes^20^ had H3K79me2 levels comparable to those MLL-AF4 targets whose expression was downregulated by pinometostat. Yet only 31% of “spreading genes” were downregulated by 10 nM pinometostat, suggesting that effects on gene expression from depletion of H3K79me2 could be governed by other factors including changes to the distribution of other chromatin modifications.

To our surprise, the pinometostat-induced activation of MHC class II genes we observed did not appear to result from a loss of H3K27me3-mediated repression, despite PRC2 subunit downregulation. Treatment with PRC2 inhibitor EI1 had no effect on *CIITA* or MHC class II gene expression but significantly reduced proliferation in MV4;11 cells (Figure 6E and S6B). A growing body of evidence supports an essential role for the PRC2 complex in MLL-r leukemogenesis--PRC2 is necessary for *MLL-AF9*-induced leukemogenesis in mouse progenitor cells and cooperates with MLL-AF9 to promote self-renewal of acute myeloid leukemia cells^82,84^. The observed downregulation of the MLL-AF4 target oncogenes upon EZH2 inhibition (Figure 6F), suggests that MLL-fusion-mediated gene activation is in some way dependent on PRC2 methyltransferase activity. Consistent with this idea, ectopic expression of *EZH2* was able to provide a small but significant proliferation rescue when treating cells with 10 nM pinometostat (Figure 6G).

We identified pinometostat-induced increases in H3K4me3 at TSS’s genome-wide (Figure 7C). Although H3K4me3 is transcriptionally activating^92,105^, the largest H3K4me3 increases were at downregulated MLL-AF4 targets that had the largest decreases in H3K79me2. Though DOT1L inhibition reduces global H3K27me3, this is unlikely to explain the massive increases in H3K4me3 that we observe^81,106^. Studies in human embryonic stem cells and mouse preadipocytes observed no genome-wide increases in H3K4me3 upon *EZH2* knockout and reductions in H3K27me3^115,116^. Rather, the increases in H3K4me3 may be due to either reduced antagonism from loss of H3K79me2 or hypermethylation due to a stalled transcriptional complex containing MLL1 near the TSS. The former scenario could function to localize H3K4me3 at the TSS, preventing spurious transcription initiation within the gene. In the latter scenario, if H3K79me2 does indeed facilitate the transition from transcription initiation to elongation^109^, its depletion could increase in H3K4me3 through greater dwelling time of RNAPII at the TSS. Indeed, given the global increases in H3K79me2 we observed upon treatment with MLL1 inhibitors (Figure S7C) it seems probable that there is a reciprocal antagonism of these two modifications on either’s deposition, potentially through H3K79me2-mediated recruitment of LSD1. Previous studies have observed that knockout or inhibition of LSD1, the H3K4me2-histone demethylase and component of the MLL-supercomplex, results in apoptosis and differentiation of MLL-r cells, inhibits leukemogenesis in mouse models and increases H3K4me2/3 at MLL-target genes^110–113^.

### Broader Clinical implications

In light of the heightened sensitivity of the FLT3-ITD lesion to DOT1L inhibition, small molecules such as pinometostat may prove effective in treating non-MLL-r leukemias with FLT3-ITD mutations. Although several FLT3 inhibitors have undergone clinical trials, drug resistance has emerged as a formidable and so far, insurmountable barrier to an effective treatment. A previous study observed that siRNAs targeting FLT3 expression increased the efficacy of the FLT3 inhibitor tandutinib^114^. As a way of circumventing the difficulties associated with siRNA delivery, DOT1L inhibitors that reduce FLT3 expression could serve as part of an effective combinatorial treatment with drugs that target FLT3 function. Our mechanistic studies provide important impetus for exploration of these ideas in pre-clinical or patient-derived FLT3-ITD leukemias.

## Acknowledgements

This work was supported by the American Cancer Society (130230-RSG-16-248-01-DMC to A.J.R.) and National Institutes of Health (R01-GM115945 to A.J.R.). We would like to thank Peter Faber and Mikayla Marchuk of the University of Chicago Functional Genomics Core and Balaji Manicassamy for his kind help with lentivirus protocols.

## Accession Information

The next-generation sequencing datasets generated and used for this study have been deposited with the Gene Expression Omnibus (GEO) under accession number GSE162441.

## Author Contributions

W.F.R. and A.J.R. conceived of and designed this study. Nearly all experiments were conducted by W.F.R with oversight by A.J.R. ICeChIP methodology used in this study was developed by R.N.S., and ICeChIP-seq was conducted by W.F.R. (all datasets except AR15) and R.N.S. (AR15 datasets). Bioinformatic analyses were conducted by W.F.R. and R.N.S. Funding acquisition was conducted by A.J.R.

## Declaration of Interests

The authors declare no competing interests.

## Materials and Methods

### Cell Culture

Human MV4;11 and MOLM13 leukemia cells and MLL1 inhibitor MM-401 were gifts from the laboratory of Yali Dou at the University of Michigan. Human THP-1 leukemia cells (cat # TIB-202) were purchased from American Type Culture Collection (ATCC). Human SEM leukemia cells (ACC546) were obtained from DSMZ-the German Collection of Microorganisms and Cell Cultures GmbH. Cells were cultured in RPMI-1640 medium containing 10% (v/v) FBessence (Seradigm cat # 3100-500), 1% L-glutamine at 37°C in humidified air containing 5% CO_2_. DOT1L inhibitor pinometostat (EPZ5676, Cayman Chemical cat # 16175), EZH2 inhibitor EI1 (Cayman Chemical cat # 19146-1), FLT3 inhibitor tandutinib (MLN518) (Selleckchem cat # S1043), MI-503 (Selleckchem cat # S7817) and PIM1 inhibitor Quercetagenin (MedChemExpress cat # HY-15604) were resuspended in DMSO. Doxycycline (Alfa Aesar cat # J60422) was resuspended in water.

### Plasmid generation

pCMV-Gag-Pol plasmid, encoding HIV-1 derived *gag*, and *pol,* the pCMV-VSV-G vector encoding VSV-G envelope gene, pTRIPZ-EZH2 and Tet-pLKO were purchased from Addgene. pTRIPZ-STAT5a-CA and pTRIPZ-FLT3-ITD were created by cloning *STAT5A* and *FLT3-ITD* from cDNA from MV4;11 cells. STAT5A-CA mutations were introduced at H298R and S710F and genes were inserted into the pTRIPZ plasmid at restriction sites AgeI and MluI. shRNA constructs were created by inserting annealed oligos of shRNA sequences (Supplementary Table I) purchased from IDT into Tet-pLKO at the AgeI and EcoRI restriction sites.

### RNA-seq and Gene Expression Analysis

Exponentially growing MV4;11 cells were grown in 150 mm^2^ tissue culture-treated plates (Corning cat # 0877224) in 30 ml media ± 10 nM pinometostat for 7 days. Every 2 days cells were spun down at 500 × g 5 min. then resuspended in media ± 10 nM pinometostat. On day 7 1 × 10^7^ cells were spun down at 500 × g 5 min. then cells were resuspended in 1 ml Trizol reagent (Life Technologies cat# 15596018), incubated 5 min. at RT then 200 μl chloroform was added and samples were shaken rigorously for 15 sec. then incubated 3 min. at RT and spun down 12,000 × g 15 min. at 4 °C. The aqueous layer (~ 500 μl) was removed and mixed with 500 μl EtOH and added to a Zymo Research RNA Clean and Concentrator column (cat # 11-353B) and spun 12,000 × g 1 min.. 100 μl DNase I (1:10 in buffered dH20) (Thermo Fisher Scientific cat # en0521) was added to the column and then spun 500 × g 5 min., incubated 15 min. at RT and then spin 12,000 × g for 30 sec. Combined 200 μl RNA binding buffer with 300 μl EtOH and then spun 12,000 × g for 30 sec. and the flow through was discarded. After each of the following were added to the column, it was spun down 12,000 × g for 30 sec. and the flow through was discarded: 400 μl RNA prep buffer; 700 μl RNA wash buffer; and 400 μl RNA wash buffer.

RNA was eluted from column with 30 μl RNase-free dH2O. Added RNA standards to 2 μg of each RNA sample-Add the equivalent of 10 copies/cell yeast RAD51; 30 copies/cell RNL2; 200 copies/cell E coli MBP; and 2000 copies/cell yeast SUMO to each sample then proceed with rRNA removal Ribo Zero Gold kit (Illumina cat # MRZ11124C) according to manufacturer’s protocol. Libraries were prepared using the NEBNext Ultra Directional RNA Library prep kit (NEB cat # E7420S). Libraries were then sequenced on the Illumina NextSEQ500. Reads were aligned to the hg38 genome assembly using HISAT2^115^ and differential gene expression analysis was done with Cufflinks^116^.

### Reverse Transcription and Quantitative real-time PCR

RNA was extracted from 10^6^ cells using 500 μl Trizol and following the manufacturer’s protocol. 1 μg RNA was used for reverse transcription with 0.5 μl MMLV HP reverse transcriptase (Lucigen cat # RT80125K) per 20 μl rxn. RNA was then degraded by alkaline hydrolysis by adding 40 μl 150 mM KOH, 20 mM tris base and heating 95 °C 10 min. then cooling on ice and quenching with 40 μl 150 mM HCl and then adding 100 μl TE. Gene expression was assayed by real-time PCR in 10 μl reactions with 0.5 μl cDNA and 5 μl PowerUP SYBR Green master mix (Applied Biosystems cat # A25742) per reactions. qPCR was run on the Bio-Rad thermocycler CFX96 or CFX384 using the program: 50 °C 2:00, 95 °C 2:00, then 40 cycles 95 °C 0:15, then 60 °C 1:00. Expression was normalized to 18S rRNA. Primer sets are listed in Supplementary Table I.

### Cell Proliferation Assay

Cells were seeded at 10^5^ cells/ml in 80 μl in clear bottom 96-well plates (Corning 07200566) in 3 replicates. Everyday 40 μl of culture was transferred to 40 μl media in a new plate. On odd days 30 μl of Cell TiterGlo 2 (Promega cat # G924A) was added to the remaining 40 μl culture and incubated 10 min. at room temperature on a shaker at 600 rpm. Luminescence was measured on a Tecan Infinite F200 Pro plate reader and fraction viability was determined from the luminescence of treated over untreated cells.

### Apoptosis Assay

Exponentially growing cells were incubated with increasing concentrations of pinometostat for 7 days in 3ml media in 6-well plates in 3 experimental replicates. 10^6^ were harvested from each plate and washed twice in 1 ml PBS then resuspended in 1 ml binding buffer as per BD Biosciences manufacturer’s protocol. Add 5 μl FITC-conjugated Annexin V (BD Biosciences cat# 556420) and 2 μl propidium iodide (Alfa Aesar cat # J66584) to 100 μl cells and incubate 15 min. at RT in the dark. Cells were then sorted on the BD FACSAriaII device for propidium iodide or FITC (Annexin V) positive cells. Data was analyzed using FlowJo software (Tree Star).

### Calibrated chromatin immunoprecipitation sequencing (ICeChIP-seq)

Native, internally calibrated ICeChIP-seq was carried out as described previously for H3K4me3 and H3K27me3^37,38^. A modified protocol was used for H3K79me2 that included cross-linking and denaturation, because of greater difficulty in immunoprecipitation of this modification, likely due to reduced accessibility of this mark within the more highly-structured nucleosome core. Briefly, nuclei were obtained from 20 million MV4;11 cells and processed to obtain HAP-purified mononucleosomal chromatin. 150 μl of 280 μl total HAP-purified chromatin was removed for denaturative ICeChIP and crosslinked in 0.25% formaldehyde for 8 min. on a nutator at RT, then quenched by adding 1M Tris pH 7.5 to 200 mM and incubating 5 min. at RT on a nutator. 50 μl of cross-linked chromatin was used for denaturation and 2.5 ul 20% SDS was added to 1% SDS final concentration and sample was incubated 1 min. at 55 °C, then IMMEDIATELY put on ice. This was then diluted with 9 volumes water (450 ul) and 100 μl was used for each IP. Antibodies for both the DMSO- and pinometostat-treated samples were processed together (12 μl antibody-bound beads per IP). 3 μg of anti-H3K79me2 (Abcam cat # ab3594, lot # GR173874); 3 μg of anti-H3K4me3 (Abcam cat # 12209, lot # GR275790-1); and 0.6 μg of anti-H3K27me3 (Cell Signaling cat # 9733, lot # 8) were used per IP. For crosslinked IPs include 1 hour 65 °C after proteinase K digest. Libraries were prepared using the NEBNext Ultra II DNA Library prep kit (NEB cat # E7645). 3 cycles of PCR amplification were used for native inputs, 4 cycles for denaturated inputs and H3K27me3 IPs, 7 cycles for H3K4me3 IPs and 10 cycles for H3K79me2 IPs. Analysis of histone methylation density (HMD) was carried out using the scripts and workflow from Grzybowski et al.^38^

### Western blotting

10 ul whole cell extracts of 2 × 10^5^ cells in 40 μl 6X SDS loading buffer were run on 4-14% bis-tris gel (Invitrogen cat # NP0335). Membranes were transferred by semi-dry apparatus (Bio-Rad Transblot cat # 170-3940) at 200 mA, 25 V for 35 min to 0.45 μm nitrocellulose membrane (Millipore cat # IPVH00010). Membranes were then blocked for 1 h with TBS-T 1% ECL Prime blocking reagent (GE Healthcare cat # RPN418) at RT on an orbital shaker and blotted with primary antibody for 1 h at RT with gentle agitation. Membranes were then washed 3 times for 5 min. while shaking with TBS-T and then incubated with secondary antibody at RT for 1 h while shaking. A complete list of antibodies can be found in Supplementary Table II.

### Transfection for lentiviral particle generation

Lentiviral particles were produced by Fugene transfection of the 293T packaging cell line in a 6-well plate at ~70% confluency with pCMV-Gag-Pol, pCMV-VSV-G and 2 μg of the plasmid encoding the gene or shRNA of interest using a 3:1:4 ratio respectively. Lentiviral particle enriched supernatants were collected 72 hours after transfection for immediate transduction.

### Lentiviral transduction

4 × 10^5^ MV4;11 cells suspended in 1 ml RPMI-1640 medium containing 10% FBessence in a 6-well plate were transduced by adding 2.5 ml of 0.45 μm filtered viral supernatants from 293T cells. Then 0.8 μl polybrene (EMD Millipore cat. # TR-1003-G)/ml transduction reagent was added to the media and the plates were wrapped with parafilm and spun down at 2000 rpm for 2 hours at room temperature then incubated O/N at 37°C in humidified air containing 5% CO_2_. After 12 hours cells were spun down and resuspended in RPMI-1640 10% FBessence. After 24 h 0.5 μg/ml puromycin was added to the wells and this selection media was refreshed every 3 days to select for transduced cells. Individual clones were purified by diluting cell cultures to 1 cell/100 μl and then plating 100 μl aliquots in a 96-well plate. Wells were visually assessed for individual clones and then grown out.

## Supplementary Figure Legends

**Figure S1.**
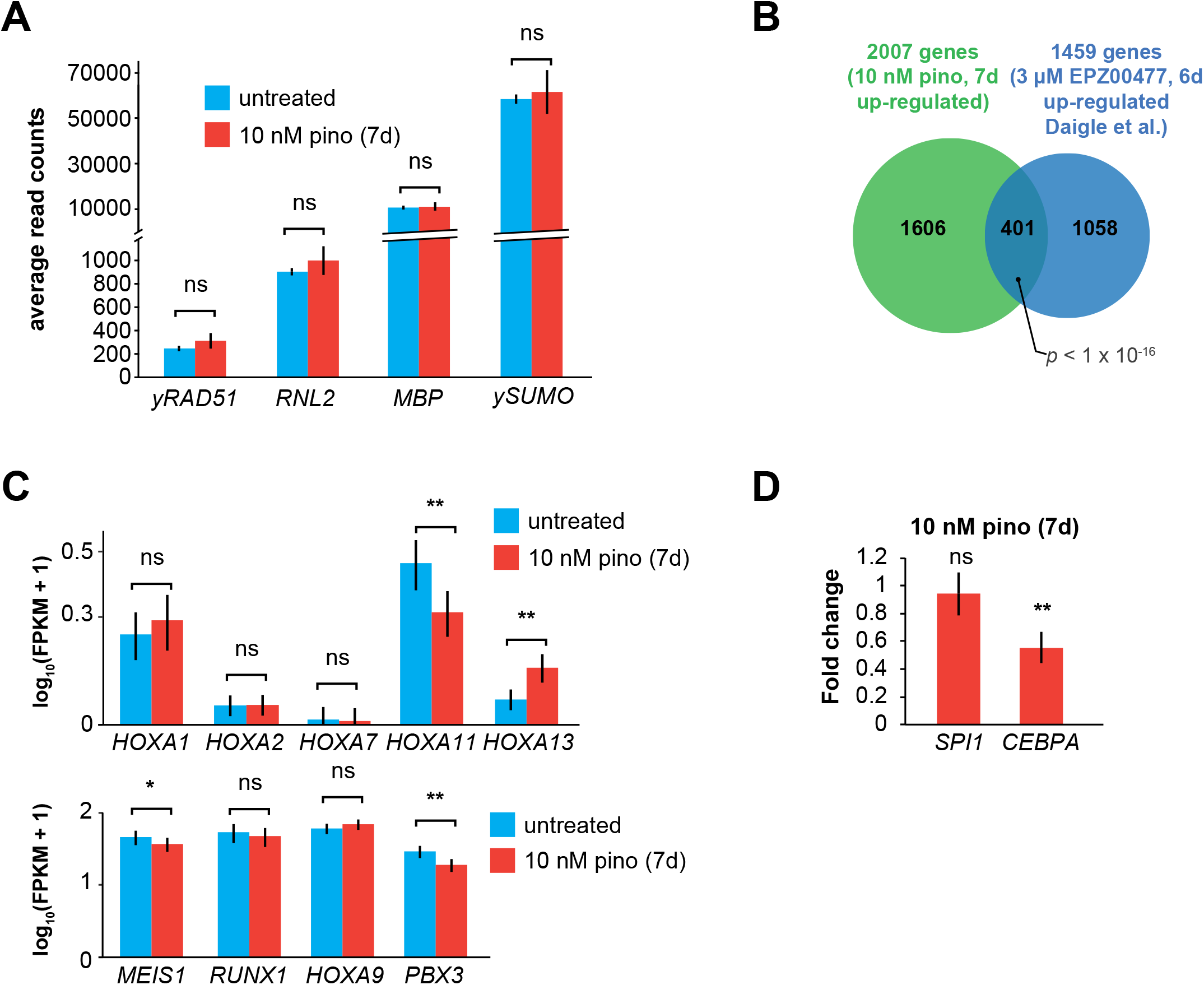
**A.** RT-qPCR analysis of non-native RNA “spike-ins” for ± 10 nM pinometostat cDNA libraries used for RNA-seq from MV4;11 cells. Bar graphs represent the average of 3 independent experiments ± S.E.M. Student’s t-test (ns *p* > 0.05). **B.** Venn diagram displaying the overlap between genes upregulated in MV4;11 cells by 10 nM pinometostat treatment (7 days) and treatment with 3 μM of the pinometostat-related compound EPZ004777 for 6 days^33^. **C.** Bar graph of Cuffdiff^116^ output for the expression of HOXA cluster genes and HOX domain-containing oncogenes *MEIS1*, *RUNX1* and *PBX3* from RNA-seq in MV4;11 cells ± 10 nM pinometostat. Values are represented as log(10) FPKM + 1 for 3 independent experiments with standard deviation. Student’s t-test (ns *p* > 0.05, * *p* < 0.05, ** *p* < 0.01). **D.** RT-qPCR analysis of *SPI1* and *CEBPA* expression from three independent experiments of MV4;11 cells treated with 10 nM pinometostat for 7 days. Fold change over DMSO-treated cells is depicted ± S.E.M. Student’s t-test (ns *p* > 0.05, ** *p* < 0.01).

**Figure S2.**
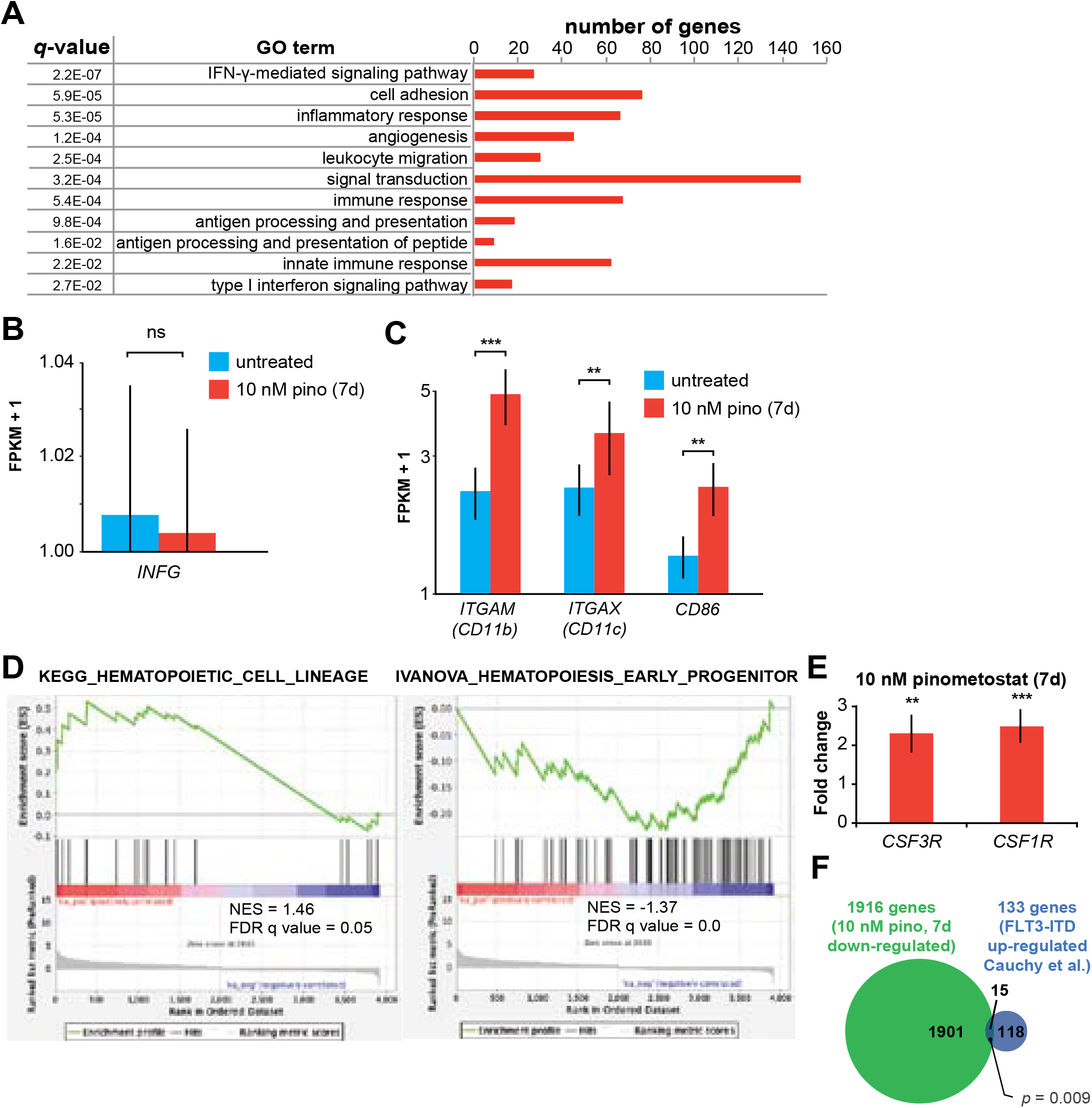
**A.** Gene Ontology analysis (DAVID)^41,42^ of pinometostat-upregulated genes showing top functional classification categories and the number of genes in each category that are significantly upregulated. **B.** Bar graph of Cuffdiff^116^ output for the expression of *INFG* (IFN-γ) from RNA-seq in MV4;11 cells ± 10 nM pinometostat. Values are represented as FPKM + 1 for 3 independent experiments with standard deviation. Student’s t-test (ns *p* > 0.05). **C.** Bar graph of Cuffdiff^116^ output for the expression of *ITGAM (CD11b), ITGAX (CD11c)* and *CD86* macrophage cell surface marker expression in MV4;11 cells ± 10 nM pinometostat for 7 days. Values are represented as FPKM + 1 for 3 independent experiments with standard deviation. Student’s t-test (** *p* < 0.01, *** *p* ≤ 0.0001). **D.** GSEA^46,47^ of the set of differentially expressed genes in MV4;11 cells ± 10 nM pinometostat compared to KEGG_HEM TOPOIETIC_CELL_LINEAGE and IVANOVA_HEMATOPOIESIS_CELL_LINEAGE gene sets from the MSigDB data base. NES-normalized enrichment score. **E.** RT-qPCR analysis of *CSF3R* and *CSF1R* expression in MV4;11 cells ± 10 nM pinometostat for 7 days. Results are displayed as mean fold-change vs. DMSO-treated cells ± S.E.M. of three independent experiments. Student’s t-test (** *p* < 0.01, *** *p* ≤ 0.0001). **F.** Venn diagram displaying the overlap between genes downregulated in MV4;11 cells by 10 nM pinometostat treatment (7 days) and genes upregulated in leukemic cells from patients with *FLT3-ITD* vs normal *FLT3* karyotypically normal AML^61^. *p*-value computed by two-tailed Fisher Exact test.

**Figure S3.**
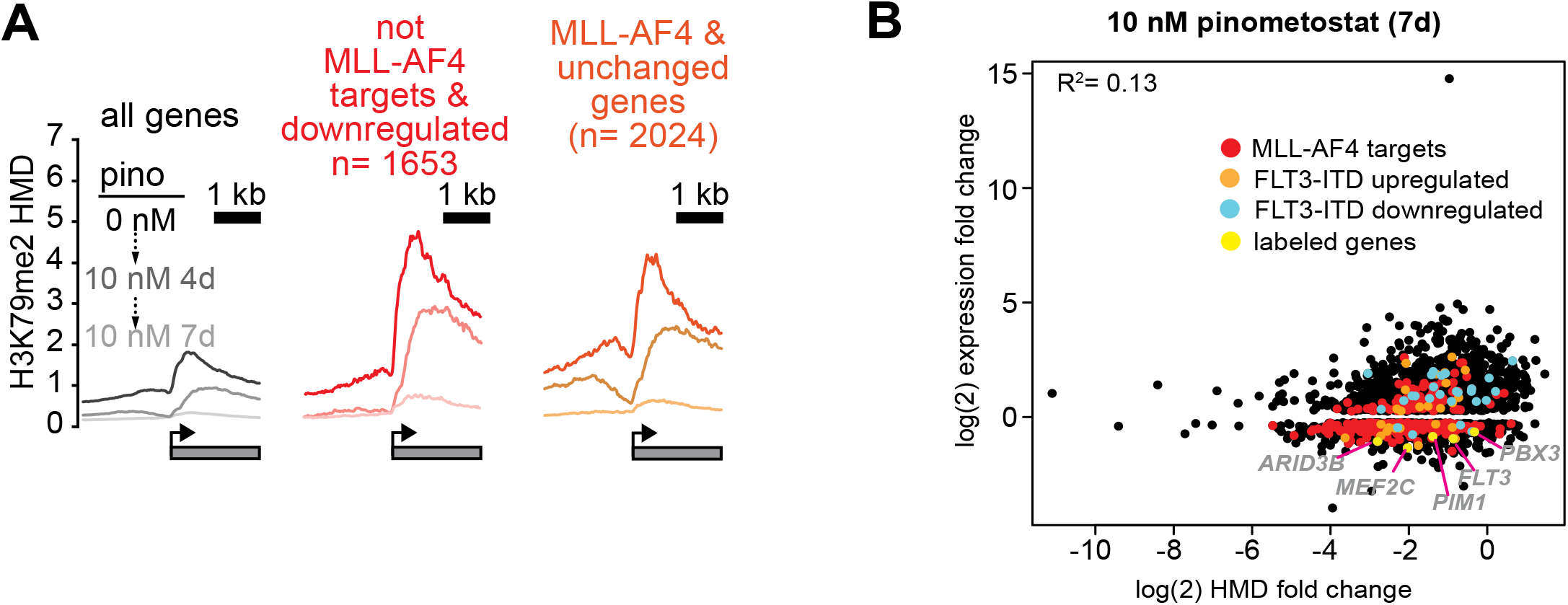
**A.** Plots of H3K79me2 density from ICeChIP-seq in MV4;11 cells treated with 10 nM pinometostat for 4 or 7 days. H3K79me2 modification density (HMD) is displayed from ± 2000 bp of the TSS of all genes expressed, non-MLL-AF4 targets^20^ downregulated by 10 nM pinometostat and MLL-AF4 targets not downregulated by 10 nM pinometostat. **B.** Scatterplot of genes downregulated by 10 nM pinometostat plotted as the log(2) fold-change in gene expression vs the log(2) fold-change in HMD with MLL-AF4 targets in red, FLT3-ITD upregulated genes^61^ in orange, FLT3-ITD downregulated genes^61^ in cyan and labeled genes highlighted in yellow. Regression analysis of all genes (R^2^ = 0.13) or the MLL-AF4 target subset (R^2^ = 0.20), reveals poor correlation between log fold changes of H3K79me2 and RNA expression.

**Figure S4.**
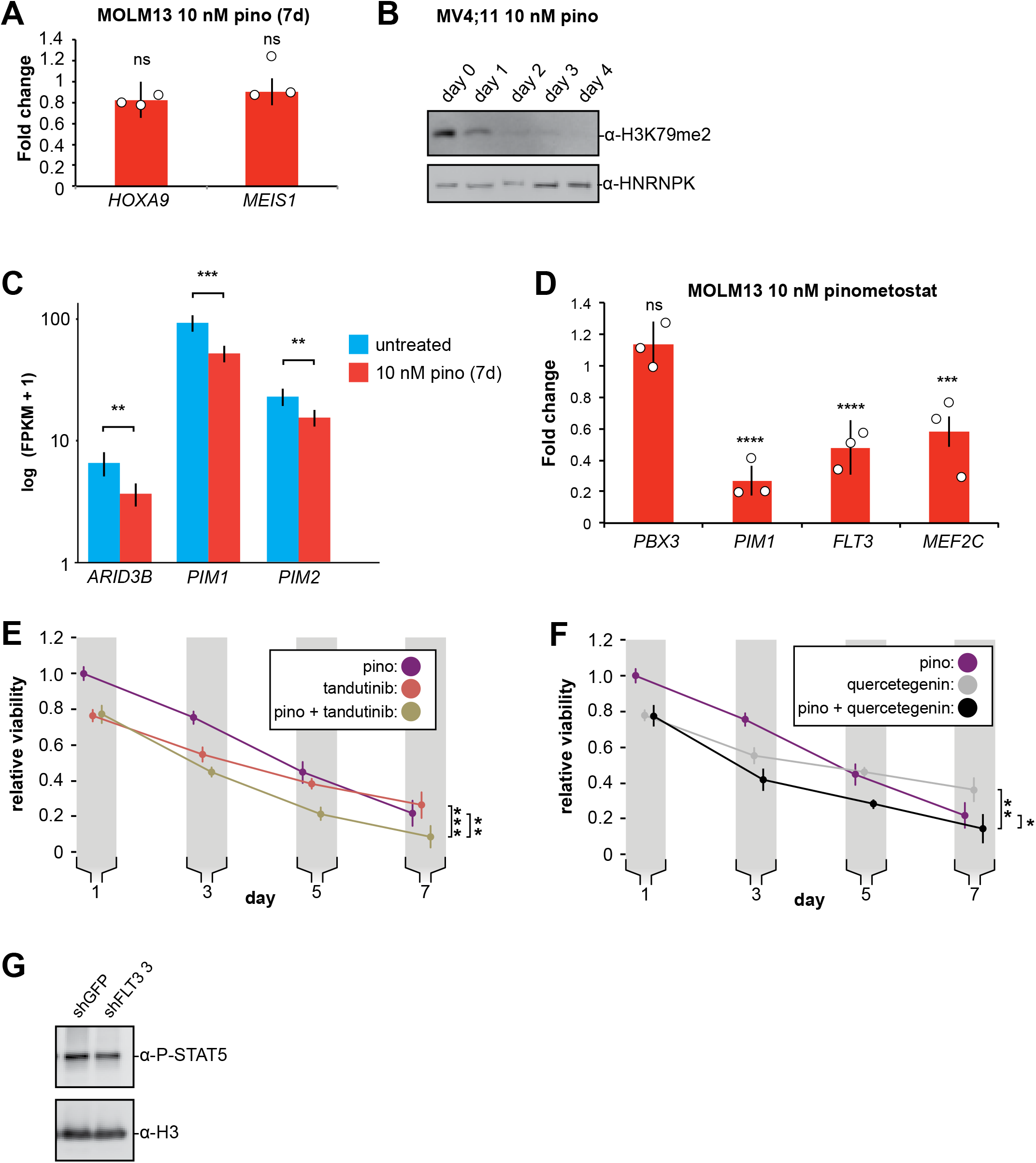
**A.** RT-qPCR analysis of *HOXA9* and *MEIS1* expression from three independent experiments of MOLM13 cells treated with 10 nM pinometostat for 7 days. Fold change over DMSO-treated cells is depicted ± S.E.M. Student’s t-test (ns *p* > 0.05). **B.** Western blots of cell extract from MV4;11 cells treated with 10 nM pinometostat for the indicated number of days and then blotted for H3K79me2 and HNRNPK as a loading control. **C.** Bar graph of Cuffdiff^116^ output for the expression of STAT5A targets *ARID3B*, *PIM1* and *PIM2* from RNA-seq in MV4;11 cells ± 10 nM pinometostat. Values are represented as log(10) FPKM + 1 for 3 independent experiments with standard deviation. Student’s t-test (** *p* < 0.01, *** *p* < 0.001). **D.** RT-qPCR analysis of *PBX3*, *PIM1, FLT3* and *MEF2C* expression from three independent experiments of MOLM13 cells treated with 10 nM pinometostat for 7 days. Fold change over DMSO-treated cells is depicted ± S.E.M. Student’s t-test (ns *p* > 0.05, *** *p* < 0.001, **** *p* < 0.0001). **E.** Proliferation assay of MV4;11 cells treated with DOT1L or FLT3 inhibitors alone or in combination using CellTiter Glo 2.0 to measure viability, showing the luminescence fraction of inhibited over uninhibited cells. Data are represented as mean ± SE of three independent experiments. Student’s t-test for significance of day 7 values: 10 nM pinometostat vs. combined ** p < 0.01, 30 nM tandutinib vs combined *** p < 0.001. **F.** Same as E but cells were treated with DOT1L and PIM1 inhibitors alone or in combination. Student’s t-test for significance of day 7 values: 10 nM pinometostat vs. combined * *p* < 0.05, 10 μM quercetagenin vs combined ** *p* < 0.01. **G.** Western blots of MV4;11 cell extract from clonal cell lines expressing shRNA to FLT3 (clone 3) or GFP blotted for phosphorylated STAT5 or histone H3 as a loading control.

**Figure S5.**
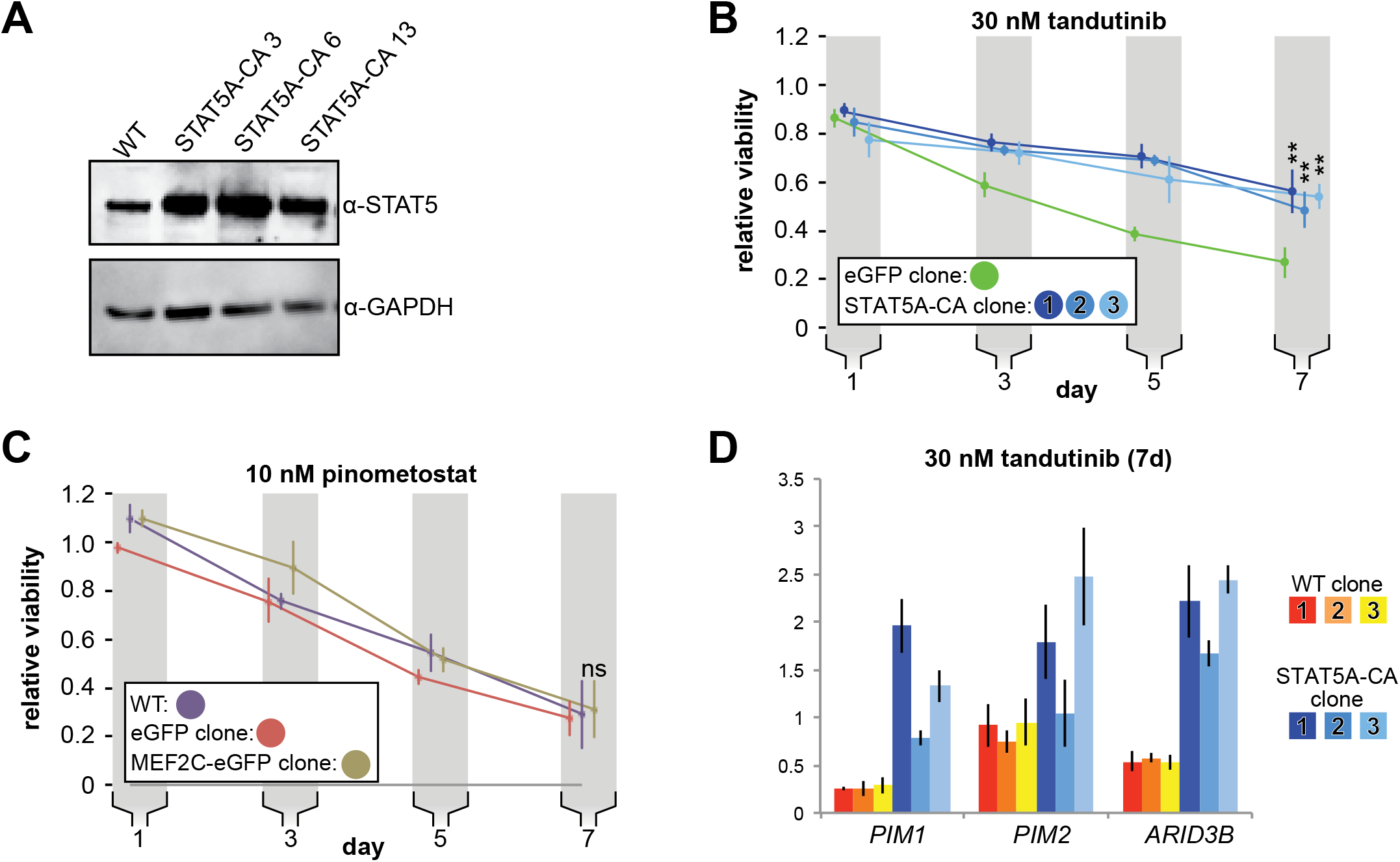
**A.** Western blots of whole cell extracts from WT MV4;11 cells or 3 monoclonal populations isolated from MV4;11 cells virally transduced with constitutively active *STAT5A* (*STAT5A-CA*). Membranes were blotted for STAT5 or GAPDH as a loading control. **B.** Proliferation assay of MV4;11 clonal isolates overexpressing *STAT5A-CA* or GFP through induction with 1 μg/mL doxycycline and treated with 30 nM tandutinib using CellTiter Glo 2.0 to measure viability, showing the luminescence fraction of inhibited over uninhibited cells, both with induced transgene (either STAT5A-CA or eGFP). Means ± SE are shown for 3 independent experiments. Student’s t-test of day 7 values: ** *p* < 0.01. **C.** Proliferation assay done as in B. with MV4;11 WT or virally transduced with *GFP* or *MEF2C-GFP* and induced to express either construct with 1 μg/mL doxycycline and treated with 10 nM pinometostat. Means #x00B1; SE are shown for 3 independent experiments. Student’s t-test of day 7 values: ns *p* > 0.05. **D.** Gene expression analysis by RT-qPCR of STAT5A target genes in WT MV4;11 cells or MV4;11 *STAT5A-CA* clones from A. induced with 1 μg/mL doxycycline to express STAT5A-CA and treated with 30 nM tandutinib for 7 days. Results are displayed as fold-change over DMSO-treated WT cells.

**Figure S6.**
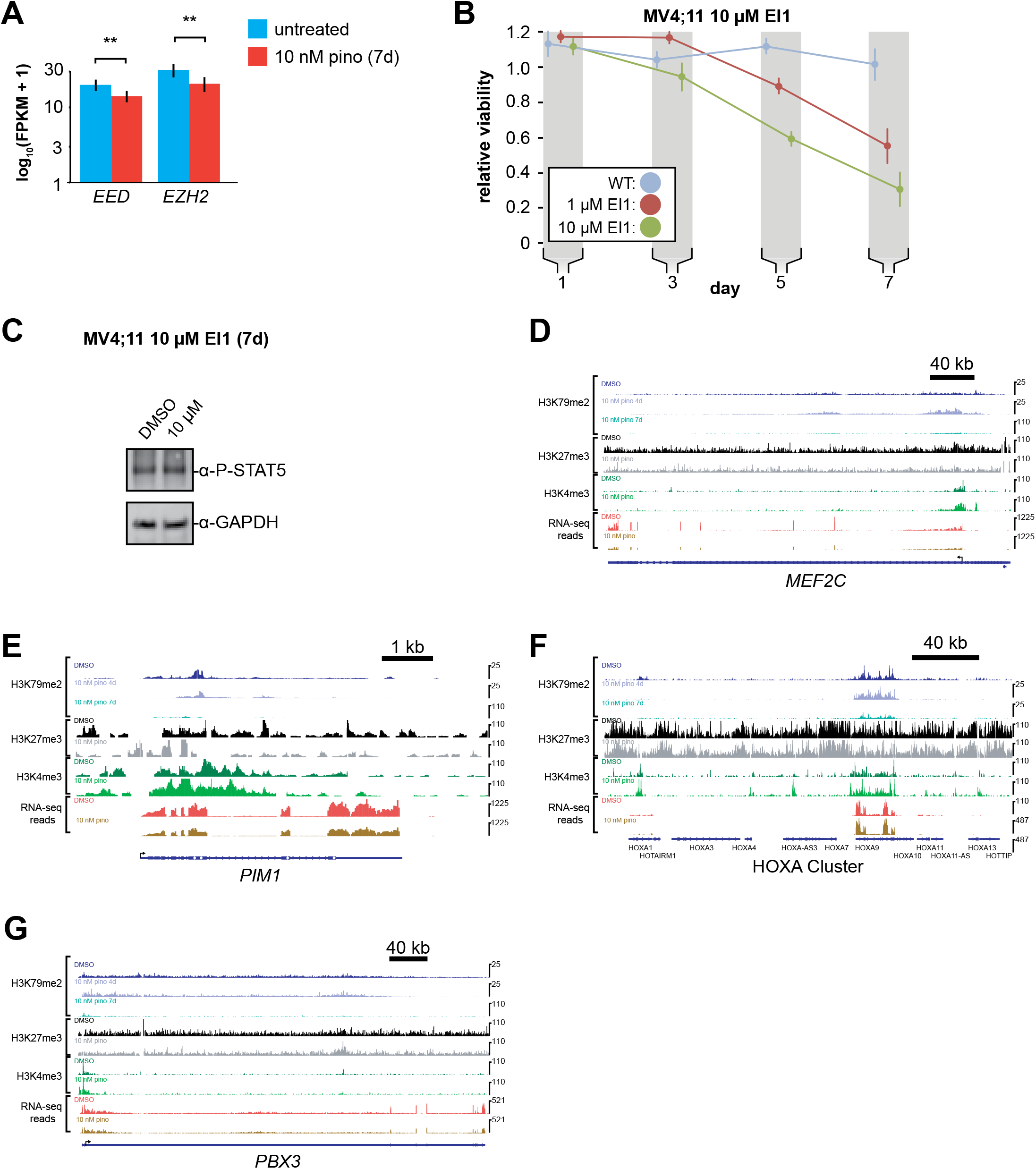
**A.** Bar graph of Cuffdiff^116^ output for the expression of polycomb complex members *EED* and *EZH2* from RNA-seq in MV4;11 cells ± 10 nM pinometostat for 7 days. Values are represented as log(10) FPKM + 1 for 3 independent experiments with standard deviation. Student’s t-test (** *p* < 0.01). **B.** Proliferation assay of MV4;11 cells treated with 10 μM EI1 EZH2 inhibitor using CellTiter Glo 2.0 to measure viability, showing the luminescence fraction of inhibited over DMSO-treated cells. Means ± SE are shown for 3 independent experiments. Student’s t-test of day 7 values: * *p* < 0.05, ** *p* < 0.01. **C.** Western blot of extract from MV4;11 cells treated with 10 μM EI1 for 7 days and blotted for phosphorylated STAT5 or GAPDH as a loading control. **D-G.** ICeChIP-seq tracks of H3K79me2, H3K27me3 and H3K4me3 HMD and an RNA-seq track (FPKM) from a single replicate for MV4;11 cells treated with 10 nM pinometostat or DMSO for 7 days for MLL-AF4^20^ and STAT5A^80^ target genes: **D.** *MEF2C*; **E.** the HOXA gene cluster; **F.** *PIM1* and **G.** *PBX3*.

**Figure S7.**
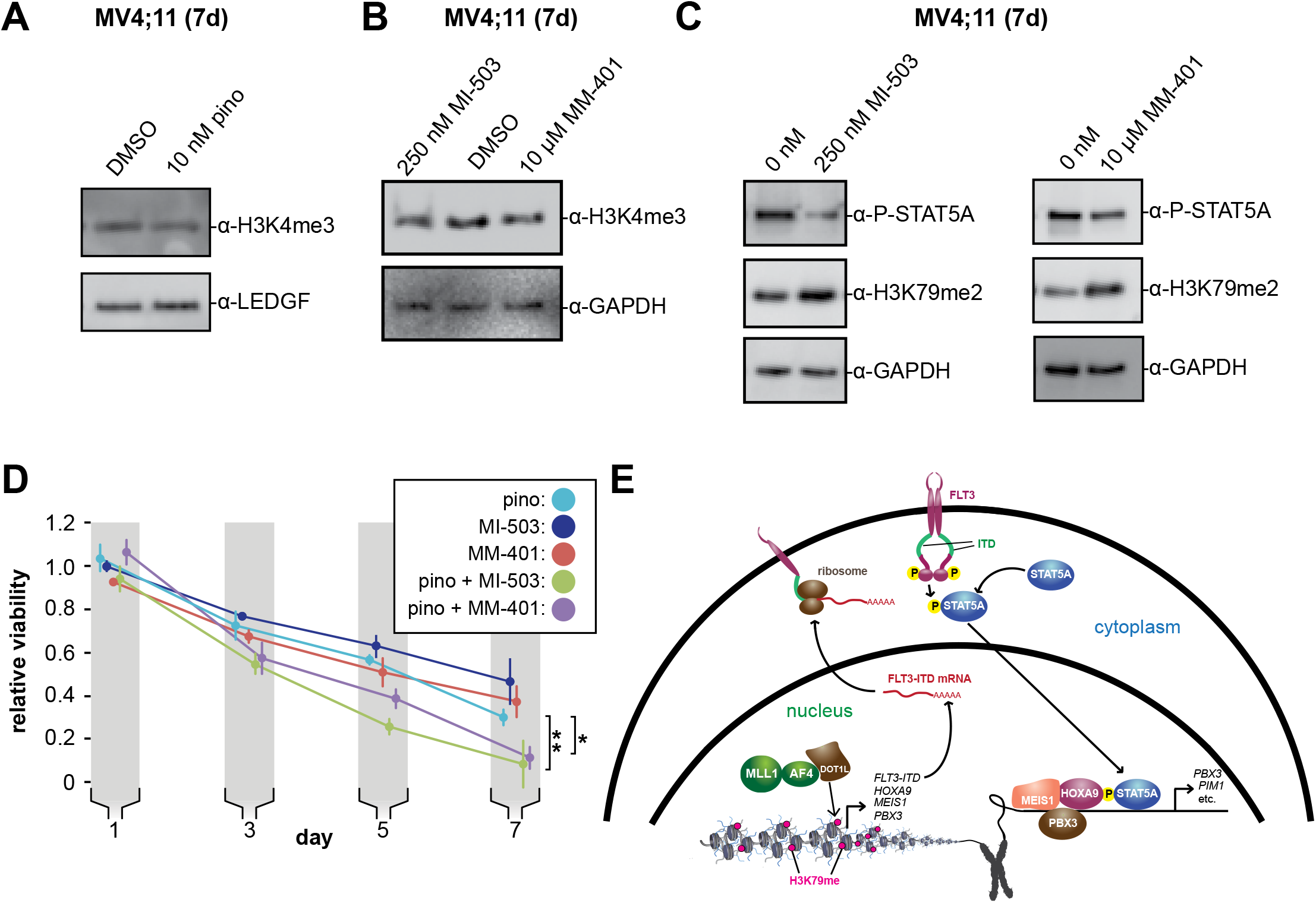
**A.** Western blots of whole cell extract from MV4;11 cells treated with 10 nM pinometostat for 7 days and blotted for H3K4me3 or LEDGF as a loading control. **B.** Western blots of whole cell extract from MV4;11 cells treated with MLL1 inhibitors MI-503 (250 nM) or MM-401 (10 μM) or DMSO for 7 days and blotted for histone H3 lysine 4 trimethylation (H3K4me3) or GAPDH as a loading control. **C.** Western blots of MV4;11 cell extract treated with MLL1 inhibitors MI-503 (250 nM) or MM-401 (10 μM) for 7 days and blotted for phosphorylated STAT5, histone H3 lysine 79 dimethylation (H3K79me2) or GAPDH as a loading control. **D.** MV4;11 cells were treated with DOT1L or MLL1 inhibitors alone or in combination for 7 days. Viability was analyzed using CellTiter Glo 2.0, showing the luminescence fraction of inhibited over DMSO-treated cells. Means ± SE are shown for 3 independent experiments. Student’s t-test of day 7 values: 10 nM pinometostat vs. 10 nM pinometostat + 250 nM MI-503 ** *p* < 0.01 10 nM pinometostat vs. 10 nM pinometostat + 10 μM MM-401 * *p* < 0.05). **E.** Model of MLL-fusion-mediated activation of HOXA9/MEIS1 and STAT5A co-targets in MLL-r, *FLT3-ITD+* leukemia. MLL-AF4 activates *HOXA9*, *MEIS1* and *FLT3-ITD* gene expression through recruitment of DOT1L and H3K79me2 hypermethylation (fuchsia). FLT3-ITD phosphorylates STAT5A allowing it to translocate to the nucleus to cooperatively bind HOXA9/MEIS1 targets with PBX3 and facilitate gene activation.

## References

1. Marks DI, Moorman A V., Chilton L, et al. The clinical characteristics, therapy and outcome of 85 adults with acute lymphoblastic leukemia and t(4;11)(q21;q23)/MLL-AFF1 prospectively treated in the UKALLXII/ECOG2993 trial. Haematologica. 2013;98(6):945–952. doi:10.3324/haematol.2012.081877

2. Jabbour E, O’Brien S, Konopleva M, Kantarjian H. New insights into the pathophysiology and therapy of adult acute lymphoblastic leukemia. Cancer. 2015;121(15):2517–2528. doi:10.1002/cncr.29383

3. Mann G, Attarbaschi A, Schrappe M, et al. Improved outcome with hematopoietic stem cell transplantation in a poor prognostic subgroup of infants with mixed-lineage-leukemia (MLL)-rearranged acute lymphoblastic leukemia: Results from the Interfant-99 Study. Blood. 2010;116(15):2644–2650. doi:10.1182/blood-2010-03-273532

4. Pieters R, Schrappe M, De Lorenzo P, et al. A treatment protocol for infants younger than 1 year with acute lymphoblastic leukaemia (Interfant-99): an observational study and a multicentre randomised trial. Lancet. 2007;370(9583):240–250. doi:10.1016/S0140-6736(07)61126-X

5. Winters AC, Bernt KM. MLL-rearranged leukemias-An update on science and clinical approaches. Front Pediatr. 2017;5(February):11–13. doi:10.3389/fped.2017.00004

6. Grossmann V, Schnittger S, Poetzinger F, et al. High incidence of RAS signalling pathway mutations in MLL-rearranged acute myeloid leukemia. Leukemia. 2013;27(9):1933–1936. doi:10.1038/leu.2013.90

7. Liang DC, Shih LY, Fu JF, et al. K-Ras mutations and N-Ras mutations in childhood acute leukemias with or without mixed-lineage leukemia gene rearrangements. Cancer. 2006;106(4):950–956. doi:10.1002/cncr.21687

8. Armstrong SA, Kung AL, Mabon ME, et al. Inhibition of FLT3 in MLL: Validation of a therapeutic target identified by gene expression based classification. Cancer Cell. 2003;3(2):173–183. doi:10.1016/S1535-6108(03)00003-5

9. Ono R, Nakajima H, Ozaki K, et al. Dimerization of MLL fusion proteins and FLT3 activation synergize to induce multiple-lineage leukemogenesis. J Clin Invest. 2005;115(4):919–929. doi:10.1172/JCI200522725

10. Corral J, Lavenir I, Impey H, et al. An MII-AF9 fusion gene made by homologous recombination causes acute leukemia in chimeric mice: A method to create fusion oncogenes. Cell. 1996;85(6):853–861. doi:10.1016/S0092-8674(00)81269-6

11. Forster A, Pannell R, Drynan LF, et al. Engineering de novo reciprocal chromosomal translocations associated with Mll to replicate primary events of human cancer. Cancer Cell. 2003;3(5):449–458. doi:10.1016/S1535-6108(03)00106-5

12. Hess JL. MLL: A histone methyltransferase disrupted in leukemia. Trends Mol Med. 2004;10(10):500–507. doi:10.1016/j.molmed.2004.08.005

13. Meyer C, Burmeister T, Gröger D, et al. The MLL recombinome of acute leukemias in 2017. Leukemia. 2018;32(2):273–284. doi:10.1038/leu.2017.213

14. Milne TA, Martin ME, Brock HW, Slany RK, Hess JL. Leukemogenic MLL fusion proteins bind across a broad region of the Hox a9 locus, promoting transcription and multiple histone modifications. Cancer Res. 2005;65(24):11367–11374. doi:10.1158/0008-5472.CAN-05-1041

15. Cao F, Townsend EC, Karatas H, et al. Targeting MLL1 H3K4 Methyltransferase Activity in Mixed-Lineage Leukemia. Mol Cell. 2014;53(2):247–261. doi:10.1016/j.molcel.2013.12.001

16. Milne TA, Kim J, Wang GG, et al. Multiple Interactions Recruit MLL1 and MLL1 Fusion Proteins to the HOXA9 Locus in Leukemogenesis. Mol Cell. 2010;38(6):853–863. doi:10.1016/j.molcel.2010.05.011

17. Marschalek R. Mechanisms of leukemogenesis by MLL fusion proteins. Br J Haematol. 2011;152(2):141–154. doi:10.1111/j.1365-2141.2010.08459.x

18. Mohan M, Herz HM, Takahashi YH, et al. Linking H3K79 trimethylation to Wnt signaling through a novel Dot1-containing complex (DotCom). Genes Dev. 2010;24(6):574–589. doi:10.1101/gad.1898410

19. Okada Y, Feng Q, Lin Y, et al. hDOT1L links histone methylation to leukemogenesis. Cell. 2005;121(2):167–178. doi:10.1016/j.cell.2005.02.020

20. Kerry J, Godfrey L, Repapi E, et al. MLL-AF4 Spreading Identifies Binding Sites that Are Distinct from Super-Enhancers and that Govern Sensitivity to DOT1L Inhibition in Leukemia. Cell Rep. 2017;18(2):482–495. doi:10.1016/j.celrep.2016.12.054

21. Bernt KM, Zhu N, Sinha AU, et al. MLL-Rearranged Leukemia Is Dependent on Aberrant H3K79 Methylation by DOT1L. Cancer Cell. 2011;20(1):66–78. doi:10.1016/j.ccr.2011.06.010

22. Guenther MG, Lawton LN, Rozovskaia T, et al. Aberrant chromatin at genes encoding stem cell regulators in human mixed-lineage leukemia. Genes Dev. 2008;22(24):3403–3408. doi:10.1101/gad.1741408

23. Stubbs MC, Kim YM, Krivtsov A V., et al. MLL-AF9 and FLT3 cooperation in acute myelogenous leukemia: Development of a model for rapid therapeutic assessment. Leukemia. 2008;22(1):66–77. doi:10.1038/sj.leu.2404951

24. Chen CW, Koche RP, Sinha AU, et al. DOT1L inhibits SIRT1-mediated epigenetic silencing to maintain leukemic gene expression in MLL-rearranged leukemia. Nat Med. 2015;21(4):335–343. doi:10.1038/nm.3832

25. Armstrong SA, Staunton JE, Silverman LB, et al. MLL translocations specify a distinct gene expression profile that distinguishes a unique leukemia. Nat Genet. 2002;30(1):41–47. doi:10.1038/ng765

26. Daigle SR, Olhava EJ, Therkelsen CA, et al. Potent inhibition of DOT1L as treatment of MLL-fusion leukemia. Blood. 2013;122(6):1017–1025. doi:10.1182/blood-2013-04-497644

27. Zeisig BB, Milne T, Garcia-Cuellar M-P, et al. Hoxa9 and Meis1 Are Key Targets for MLL-ENL-Mediated Cellular Immortalization. Mol Cell Biol. 2004;24(2):617–628. doi:10.1128/mcb.24.2.617-628.2004

28. Chang MJ, Wu H, Achille NJ, et al. Histone H3 Lysine 79 Methyltransferase Dot1 Is Required for Immortalization by MLL Oncogenes. Cancer Res. 2010;70(24):10234–10242. doi:10.1158/0008-5472.CAN-10-3294

29. Jo SY, Granowicz EM, Maillard I, Thomas D, Hess JL. Requirement for Dot1l in murine postnatal hematopoiesis and leukemogenesis by MLL translocation. Blood. 2011;117(18):4759–4768. doi:10.1182/blood-2010-12-327668

30. Kroon E, Krosl J, Thorsteinsdottir U, Baban S, Buchberg AM, Sauvageau G. Hoxa9 transforms primary bone marrow cells through specific collaboration with Meis1a but not Pbx1b. EMBO J. 1998;17(13):3714–3725. doi:10.1093/emboj/17.13.3714

31. Calvo KR, Sykes DB, Pasillas MP, Kamps MP. Nup98-Hoxa9 immortalizes myeloid progenitors, enforces expression of Hoxa9, Hoxa7 and Meis1, and alters cytokine-specific responses in a manner similar to that induced by retroviral co-expression of Hoxa9 and Meis1. Oncogene. 2002;21(27):4247–4256. doi:10.1038/sj.onc.1205516

32. Dobson CL, Warren AJ, Pannell R, et al. The Mll-AF9 gene fusion in mice controls myeloproliferation and specifies acute myeloid leukaemogenesis. EMBO J. 1999;18(13):3564–3574. doi:10.1093/emboj/18.13.3564

33. Daigle SR, Olhava EJ, Therkelsen CA, et al. Selective Killing of Mixed Lineage Leukemia Cells by a Potent Small-Molecule DOT1L Inhibitor. Cancer Cell. 2011;20(1):53–65. doi:10.1016/j.ccr.2011.06.009

34. Deshpande AJ, Chen L, Fazio M, et al. Leukemic transformation by the MLL-AF6 fusion oncogene requires the H3K79 methyltransferase Dot1l. Blood. 2013;121(13):2533–2541. doi:10.1182/blood-2012-11-465120

35. Anglin JL, Song Y. A medicinal chemistry perspective for targeting histone H3 lysine-79 methyltransferase DOT1L. J Med Chem. 2013;56(22):8972–8983. doi:10.1021/jm4007752

36. Yu W, Chory EJ, Wernimont AK, et al. Catalytic site remodelling of the DOT1L methyltransferase by selective inhibitors. Nat Commun. 2012;3:1–12. doi:10.1038/ncomms2304

37. Grzybowski AT, Chen Z, Ruthenburg AJ. Calibrating ChIP-Seq with Nucleosomal Internal Standards to Measure Histone Modification Density Genome Wide. Mol Cell. 2015;58(5):886–899. doi:10.1016/j.molcel.2015.04.022

38. Grzybowski AT, Shah RN, Richter WF, Ruthenburg AJ. Native internally calibrated chromatin immunoprecipitation for quantitative studies of histone post-translational modifications. Nat Protoc. 2019;14(12):3275–3302. doi:10.1038/s41596-019-0218-7

39. Godfrey L, Crump NT, Thorne R, et al. DOT1L inhibition reveals a distinct subset of enhancers dependent on H3K79 methylation. Nat Commun. 2019;10(1). doi:10.1038/s41467-019-10844-3

40. Okuda H, Stanojevic B, Kanai A, et al. Cooperative gene activation by AF4 and DOT1L drives MLL-rearranged leukemia. J Clin Invest. 2017;127(5):1918–1931. doi:10.1172/JCI91406

41. Huang DW, Sherman BT, Lempicki RA. Bioinformatics enrichment tools: Paths toward the comprehensive functional analysis of large gene lists. Nucleic Acids Res. 2009;37(1):1–13. doi:10.1093/nar/gkn923

42. Huang DW, Sherman BT, Lempicki RA. Systematic and integrative analysis of large gene lists using DAVID bioinformatics resources. Nat Protoc. 2009;4(1):44–57. doi:10.1038/nprot.2008.211

43. Caldarelli A, Müller JP, Paskowski-Rogacz M, et al. A genome-wide RNAi screen identifies proteins modulating aberrant FLT3-ITD signaling. Leukemia. 2013;27(12):2301–2310. doi:10.1038/leu.2013.83

44. Spiekermann K, Pau M, Schwab R, Schmieja K, Franzrahe S, Hiddemann W. Constitutive activation of STAT3 and STAT5 is induced by leukemic fusion proteins with protein tyrosine kinase activity and is sufficient for transformation of hematopoietic precursor cells. Exp Hematol. 2002;30(3):262–271. doi:10.1016/S0301-472X(01)00787-1

45. Muhlethaler-Mottet A, Berardino W Di, Otten LA, Mach B. Activation of the MHC class II transactivator CIITA by interferon-γ requires cooperative interaction between Stat1 and USF-1. Immunity. 1998;8(2):157–166. doi:10.1016/S1074-7613(00)80468-9

46. Subramanian A, Tamayo P, Mootha VK, et al. Gene set enrichment analysis: A knowledge-based approach for interpreting genome-wide expression profiles. Proc Natl Acad Sci U S A. 2005;102(43):15545–15550. doi:10.1073/pnas.0506580102

47. Mootha VK, Lindgren CM, Eriksson K-F, et al. PGC-1α-responsive genes involved in oxidative phosphorylation are coordinately downregulated in human diabetes. Nat Genet. 2003;34(3):267–273. doi:10.1038/ng1180

48. Mossadegh-Keller N, Sarrazin S, Kandalla PK, et al. M-CSF instructs myeloid lineage fate in single haematopoietic stem cells. Nature. 2013;497(7448):239–243. doi:10.1038/nature12026

49. Klimiankou M, Dannenmann B, Solovyeva A, et al. Effects of CSF3R mutations on Myeloid Differentiation and Proliferation of Hematopoietic Cells of Congenital Neutropenia Patients. Blood. 2017;130(Supplement 1):2278. doi:10.1182/blood.V130.Suppl_1.2278.2278

50. Wilkinson AC, Ballabio E, Geng H, et al. RUNX1 Is a Key Target in t(4;11) Leukemias that Contributes to Gene Activation through an AF4-MLL Complex Interaction. Cell Rep. 2013;3(1):116–127. doi:10.1016/j.celrep.2012.12.016

51. Nagel G, Weber D, Fromm E, et al. Epidemiological, genetic, and clinical characterization by age of newly diagnosed acute myeloid leukemia based on an academic population-based registry study (AMLSG BiO). Ann Hematol. 2017;96(12):1993–2003. doi:10.1007/s00277-017-3150-3

52. Krivtsov A V., Twomey D, Feng Z, et al. Transformation from committed progenitor to leukaemia stem cell initiated by MLL-AF9. Nature. 2006;442(7104):818–822. doi:10.1038/nature04980

53. Li Z, Chen P, Su R, et al. PBX3 and MEIS1 Cooperate in hematopoietic cells to drive acute myeloid leukemias characterized by a core transcriptome of the MLL-rearranged disease. Cancer Res. 2016;76(3):619–629. doi:10.1158/0008-5472.CAN-15-1566

54. Levis M, Small D. FLT3: ITDoes matter in leukemia. Leukemia. 2003;17(9):1738–1752. doi:10.1038/sj.leu.2403099

55. Mizuki M, Schwäble J, Steur C, et al. Suppression of myeloid transcription factors and induction of STAT response genes by AML-specific Flt3 mutations. Blood. 2003;101(8):3164–3173. doi:10.1182/blood-2002-06-1677

56. Levis M, Allebach J, Tse KF, et al. A FLT3-targeted tyrosine kinase inhibitor is cytotoxic to leukemia cells in vitro and in vivo. Blood. 2002;99(11):3885–3891. doi:10.1182/blood.V99.11.3885

57. Du Y, Spence SE, Jenkins NA, Copeland NG. Cooperating cancer-gene identification through oncogenic-retrovirus-induced insertional mutagenesis. Blood. 2005;106(7):2498–2505. doi:10.1182/blood-2004-12-4840

58. Li Z, Zhang Z, Li Y, et al. PBX3 is an important cofactor of HOXA9 in leukemogenesis. Blood. 2013;121(8):1422–1431. doi:10.1182/blood-2012-07-442004

59. Wang GG, Pasillas MP, Kamps MP. Persistent Transactivation by Meis1 Replaces Hox Function in Myeloid Leukemogenesis Models: Evidence for Co-Occupancy of Meis1-Pbx and Hox-Pbx Complexes on Promoters of Leukemia-Associated Genes. Mol Cell Biol. 2006;26(10):3902–3916. doi:10.1128/mcb.26.10.3902-3916.2006

60. Stehling-Sun S, Dade J, Nutt SL, DeKoter RP, Camargo FD. Regulation of lymphoid versus myeloid fate “choice” by the transcription factor Mef2c. Nat Immunol. 2009;10(3):289–296. doi:10.1038/ni.1694

61. Cauchy P, James SR, Zacarias-Cabeza J, et al. Chronic FLT3-ITD Signaling in Acute Myeloid Leukemia Is Connected to a Specific Chromatin Signature. Cell Rep. 2015;12(5):821–836. doi:10.1016/j.celrep.2015.06.069

62. Orlando DA, Chen MW, Brown VE, et al. Quantitative ChIP-Seq normalization reveals global modulation of the epigenome. Cell Rep. 2014;9(3):1163–1170. doi:10.1016/j.celrep.2014.10.018

63. Shah RN, Grzybowski AT, Cornett EM, et al. Examining the Roles of H3K4 Methylation States with Systematically Characterized Antibodies. Mol Cell. 2018;72(1):162–177.e7. doi:10.1016/j.molcel.2018.08.015

64. Schübeler D, MacAlpine DM, Scalzo D, et al. The histone modification pattern of active genes revealed through genome-wide chromatin analysis of a higher eukaryote. Genes Dev. 2004;18(11):1263–1271. doi:10.1101/gad.1198204

65. Guenther MG, Levine SS, Boyer LA, Jaenisch R, Young RA. A Chromatin Landmark and Transcription Initiation at Most Promoters in Human Cells. Cell. 2007;130(1):77–88. doi:10.1016/j.cell.2007.05.042

66. Quentmeier H, Reinhardt J, Zaborski M, Drexler HG. FLT3 mutations in acute myeloid leukemia cell lines. Leukemia. 2003;17(1):120–124. doi:10.1038/sj.leu.2402740

67. Clark JJ, Cools J, Curley DP, et al. Variable sensitivity of FLT3 activation loop mutations to the small molecule tyrosine kinase inhibitor MLN518. Blood. 2004;104(9):2867–2872. doi:10.1182/blood-2003-12-4446

68. Choudhary C, Müller-Tidow C, Berdel WE, Serve H. Signal transduction of oncogenic Flt3. Int J Hematol. 2005;82(2):93–99. doi:10.1532/IJH97.05090

69. Onishi M, Nosaka T, Misawa K, et al. Identification and Characterization of a Constitutively Active STAT5 Mutant That Promotes Cell Proliferation. Mol Cell Biol. 1998;18(7):3871–3879. doi:10.1128/mcb.18.7.3871

70. Choudhary C, Brandts C, Schwable J, et al. Activation mechanisms of STAT5 by oncogenic Flt3-ITD. Blood. 2007;110(1):370–374. doi:10.1182/blood-2006-05-024018

71. Wierenga ATJ, Vellenga E, Schuringa JJ. Maximal STAT5-Induced Proliferation and Self-Renewal at Intermediate STAT5 Activity Levels. Mol Cell Biol. 2008;28(21):6668–6680. doi:10.1128/mcb.01025-08

72. Kim KT, Baird K, Ahn JY, et al. Pim-1 is up-regulated by constitutively activated FLT3 and plays a role in FLT3-mediated cell survival. Blood. 2005;105(4):1759–1767. doi:10.1182/blood-2004-05-2006

73. Ribeiro D, Melão A, van Boxtel R, et al. STAT5 is essential for IL-7– mediated viability, growth, and proliferation of T-cell acute lymphoblastic leukemia cells. Blood Adv. 2018;2(17):2199–2213. doi:10.1182/bloodadvances.2018021063

74. Amson R, Sigaux F, Przedborski S, Flandrin G, Givol D, Telerman A. The human protooncogene product p33pim is expressed during fetal hematopoiesis and in diverse leukemias. Proc Natl Acad Sci U S A. 1989;86(22):8857–8861. doi:10.1073/pnas.86.22.8857

75. Cibull TL, Jones TD, Li L, et al. Overexpression of Pim-1 during progression of prostatic adenocarcinoma. J Clin Pathol. 2006;59(3):285–288. doi:10.1136/jcp.2005.027672

76. Peltola K, Hollmen M, Maula SM, et al. Pim-1 kinase expression predicts radiation response in squamocellular carcinoma of head and neck and is under the control of epidermal growth factor receptor. Neoplasia. 2009;11(7):629–636. doi:10.1593/neo.81038

77. Deneen B, Welford SM, Ho T, Hernandez F, Kurland I, Denny CT. PIM3 Proto-Oncogene Kinase Is a Common Transcriptional Target of Divergent EWS/ETS Oncoproteins. Mol Cell Biol. 2003;23(11):3897–3908. doi:10.1128/mcb.23.11.3897-3908.2003

78. Adam M, Pogacic V, Bendit M, et al. Targeting PIM kinases impairs survival of hematopoietic cells transformed by kinase inhibitor-sensitive and kinase inhibitor-resistant forms of Fms-like tyrosine kinase 3 and BCR/ABL. Cancer Res. 2006;66(7):3828–3835. doi:10.1158/0008-5472.CAN-05-2309

79. Green AS, Maciel TT, Hospital MA, et al. Pim kinases modulate resistance to FLT3 tyrosine kinase inhibitors in FLT3-ITD acute myeloid leukemia. Sci Adv. 2015;1(8):1–14. doi:10.1126/sciadv.1500221

80. Moore MAS, Dorn DC, Schuringa JJ, Chung KY, Morrone G. Constitutive activation of Flt3 and STAT5A enhances self-renewal and alters differentiation of hematopoietic stem cells. Exp Hematol. 2007;35(4 SUPPL.):105–116. doi:10.1016/j.exphem.2007.01.018

81. Kim D-H, Tang Z, Shimada M, et al. Histone H3K27 Trimethylation Inhibits H3 Binding and Function of SET1-Like H3K4 Methyltransferase Complexes. Mol Cell Biol. 2013;33(24):4936–4946. doi:10.1128/mcb.00601-13

82. Shi J, Wang E, Zuber J, et al. The Polycomb complex PRC2 supports aberrant self-renewal in a mouse model of MLL-AF9;Nras G12D acute myeloid leukemia. Oncogene. 2013;32(7):930–938. doi:10.1038/onc.2012.110

83. Zhou J, Bi C, Cheong LL, et al. The histone methyltransferase inhibitor, DZNep, up-regulates TXNIP, increases ROS production, and targets leukemia cells in AML. Blood. 2011;118(10):2830–2839. doi:10.1182/blood-2010-07-294827

84. Neff T, Sinha AU, Kluk MJ, et al. Polycomb repressive complex 2 is required for MLL-AF9 leukemia. Proc Natl Acad Sci U S A. 2012;109(13):5028–5033. doi:10.1073/pnas.1202258109

85. Ueda K, Yoshimi A, Kagoya Y, et al. Inhibition of histone methyltransferase EZH2 depletes leukemia stem cell of mixed lineage leukemia fusion leukemia through upregulation of p16. Cancer Sci. 2014;105(5):512–519. doi:10.1111/cas.12386

86. Borkin D, He S, Miao H, et al. Pharmacologic inhibition of the menin-MLL interaction blocks progression of MLL leukemia invivo. Cancer Cell. 2015;27(4):589–602. doi:10.1016/j.ccell.2015.02.016

87. Li BE, Gan T, Meyerson M, Rabbitts TH, Ernst P. Distinct pathways regulated by menin and by MLL1 in hematopoietic stem cells and developing B cells. Blood. 2013;122(12):2039–2046. doi:10.1182/blood-2013-03-486647

88. Yokoyama A, Somervaille TCP, Smith KS, Rozenblatt-Rosen O, Meyerson M, Cleary ML. The menin tumor suppressor protein is an essential oncogenic cofactor for MLL-associated leukemogenesis. Cell. 2005;123(2):207–218. doi:10.1016/j.cell.2005.09.025

89. Chen S, Yang Z, Wilkinson AW, et al. The PZP Domain of AF10 Senses Unmodified H3K27 to Regulate DOT1L-Mediated Methylation of H3K79. Mol Cell. 2015;60(2):319–327. doi:10.1016/j.molcel.2015.08.019

90. Schmitges FW, Prusty AB, Faty M, et al. Histone Methylation by PRC2 Is Inhibited by Active Chromatin Marks. Mol Cell. 2011;42(3):330–341. doi:10.1016/j.molcel.2011.03.025

91. Voigt P, LeRoy G, Drury WJ, et al. Asymmetrically modified nucleosomes. Cell. 2012;151(1):181–193. doi:10.1016/j.cell.2012.09.002

92. Vermeulen M, Mulder KW, Denissov S, et al. Selective Anchoring of TFIID to Nucleosomes by Trimethylation of Histone H3 Lysine 4. Cell. 2007;131(1):58–69. doi:10.1016/j.cell.2007.08.016

93. Yokoyama A, Wang Z, Wysocka J, et al. Leukemia Proto-Oncoprotein MLL Forms a SET1-Like Histone Methyltransferase Complex with Menin To Regulate Hox Gene Expression. Mol Cell Biol. 2004;24(13):5639–5649. doi:10.1128/mcb.24.13.5639-5649.2004

94. Krogan NJ, Dover J, Khorrami S, et al. COMPASS, a histone H3 (lysine 4) methyltransferase required for telomeric silencing of gene expression. J Biol Chem. 2002;277(13):10753–10755. doi:10.1074/jbc.C200023200

95. Rosen DB, Minden MD, Kornblau SM, et al. Functional characterization of FLT3 receptor signaling deregulation in acute myeloid leukemia by single cell network profiling (SCNP). PLoS One. 2010;5(10):10–18. doi:10.1371/journal.pone.0013543

96. Zhou J, Bi C, Janakakumara J V., et al. Enhanced activation of STAT pathways and overexpression of survivin confer resistance to FLT3 inhibitors and could be therapeutic targets in AML. Blood. 2009;113(17):4052–4062. doi:10.1182/blood-2008-05-156422

97. Spiekermann K, Bagrintseva K, Schwab R, Schmieja K, Hiddemann W. Overexpression and constitutive activation of FLT3 induces STAT5 activation in primary acute myeloid leukemia blast cells. Clin Cancer Res. 2003;9(6):2140–2150.

98. Santos SCR, Lacronique V, Bouchaert I, et al. Constitutively active STAT5 variants induce growth and survival of hematopoietic cells through a PI 3-kinase/Akt dependent pathway. Oncogene. 2001;20(17):2080–2090. doi:10.1038/sj.onc.1204308

99. Fathi AT, Arowojolu O, Swinnen I, et al. A potential therapeutic target for FLT3-ITD AML: PIM1 kinase. Leuk Res. 2012;36(2):224–231. doi:10.1016/j.leukres.2011.07.011

100. Huang Y, Sitwala K, Bronstein J, et al. Identification and characterization of Hoxa9 binding sites in hematopoietic cells. Blood. 2012;119(2):388–398. doi:10.1182/blood-2011-03-341081

101. de Bock CE, Demeyer S, Degryse S, et al. HOXA9 cooperates with activated JAK/STAT signaling to drive leukemia development. Cancer Discov. 2018;8(5):616–631. doi:10.1158/2159-8290.CD-17-0583

102. Zhu J, Cote-Sierra J, Guo L, Paul WE. Stat5 activation plays a critical role in Th2 differentiation. Immunity. 2003;19(5):739–748. doi:10.1016/S1074-7613(03)00292-9

103. Rani A, Murphy JJ. STAT5 in Cancer and Immunity. J Interf Cytokine Res. 2016;36(4):226–237. doi:10.1089/jir.2015.0054

104. Yamamoto M, Kato T, Hotta C, et al. Shared and distinct functions of the transcription factors IRF4 and IRF8 in myeloid cell development. PLoS One. 2011;6(10):2–11. doi:10.1371/journal.pone.0025812

105. Chang P-Y, Hom RA, Musselman CA, et al. Binding of the MLL PHD3 Finger to Histone H3K4me3 Is Required for MLL-Dependent Gene Transcription. J Mol Biol. 2010;400(2):137–144. doi:https://doi.org/10.1016/j.jmb.2010.05.005

106. Hanson RD, Hess JL, Yu BD, et al. Mammalian Trithorax and Polycomb-group homologues are antagonistic regulators of homeotic development. Proc Natl Acad Sci U S A. 1999;96(25):14372–14377. doi:10.1073/pnas.96.25.14372

107. Collinson A, Collier AJ, Morgan NP, et al. Deletion of the Polycomb-Group Protein EZH2 Leads to Compromised Self-Renewal and Differentiation Defects in Human Embryonic Stem Cells. Cell Rep. 2016;17(10):2700–2714. doi:10.1016/j.celrep.2016.11.032

108. Wang L, Jin Q, Lee JE, Su IH, Ge K. Histone H3K27 methyltransferase Ezh2 represses Wnt genes to facilitate adipogenesis. Proc Natl Acad Sci U S A. 2010;107(16):7317–7322. doi:10.1073/pnas.1000031107

109. Wood K, Tellier M, Murphy S. DOT1L and H3K79 methylation in transcription and genomic stability. Biomolecules. 2018;8(1):1–16. doi:10.3390/biom8010011

110. Harris WJ, Huang X, Lynch JT, et al. The Histone Demethylase KDM1A Sustains the Oncogenic Potential of MLL-AF9 Leukemia Stem Cells. Cancer Cell. 2012;21(4):473–487. doi:10.1016/j.ccr.2012.03.014

111. Feng Z, Yao Y, Zhou C, et al. Pharmacological inhibition of LSD1 for the treatment of MLL-rearranged leukemia. J Hematol Oncol. 2016;9(1):1–13. doi:10.1186/s13045-016-0252-7

112. McGrath JP, Williamson KE, Balasubramanian S, et al. Pharmacological inhibition of the histone lysine demethylase KDM1A suppresses the growth of multiple acute myeloid leukemia subtypes. Cancer Res. 2016;76(7):1975–1988. doi:10.1158/0008-5472.CAN-15-2333

113. Fang J, Ying H, Mao T, et al. Upregulation of CD11b and CD86 through LSD1 inhibition promotes myeloid differentiation and suppresses cell proliferation in human monocytic leukemia cells. Oncotarget. 2017;8(49):85085–85101. doi:10.18632/oncotarget.18564

114. Walters DK, Stoffregen EP, Heinrich MC, Deininger MW, Druker BJ. Brief report RNAi-induced down-regulation of FLT3 expression in AML cell lines increases sensitivity to MLN518. 2017;105(7):2952–2955. doi:10.1182/blood-2004-07-2758.Supported

115. Kim D, Langmead B, Salzberg SL. HISAT: a fast spliced aligner with low memory requirements. Nat Methods. 2015;12(4):357–360. doi:10.1038/nmeth.3317

116. Trapnell C, Roberts A, Goff L, Geo P. Differential gene and transcript expression analysis of RNA-seq experiments with TopHat and Cufflinks. Nat Protoc. 2013;7(3):562–578. doi:10.1038/nprot.2012.016.Differential

117. Yuan W, Xie J, Long C, et al. Heterogeneous nuclear ribonucleoprotein L is a subunit of human KMT3a/set2 complex required for H3 Lys-36 trimethylation activity in vivo. J Biol Chem. 2009;284(23):15701–15707. doi:10.1074/jbc.M808431200

118. Scheeren FA, Naspetti M, Diehl S, et al. STAT5 regulates the self-renewal capacity and differentiation of human memory B cells and controls Bcl-6 expression. Nat Immunol. 2005;6(3):303–313. doi:10.1038/ni1172

